# All exons are not created equal - Exon vulnerability determines the effect of exonic mutations on splicing

**DOI:** 10.1101/2023.06.14.544306

**Authors:** Lise L. Holm, Thomas K. Doktor, Katharina K. Flugt, Ulrika S. S. Petersen, Rikke Pedersen, Brage S. Andresen

## Abstract

It is now widely accepted that aberrant splicing of constitutive exons is often caused by mutations affecting *cis-*acting splicing regulatory elements (SREs), but there is a misconception that all exons have an equal dependency on SREs and thus a similar vulnerability to aberrant splicing. We demonstrate that some exons are more likely to be affected by exonic splicing mutations (ESM) due to an inherent vulnerability, which is context-dependent and influenced by the strength of exon definition. We have developed VulExMap, a tool which based on empirical data that can designate whether a constitutive exon is vulnerable. Using VulExMap, we find that only 27% of all exons can be categorized as vulnerable whereas two-thirds of 332 previously reported ESMs in 71 disease genes are located in vulnerable exons. Because VulExMap analysis is based on empirical data on splicing of exons in their endogenous context, it includes all features important in determining the vulnerability. We believe that VulExMap will be an important tool when assessing the effect of exonic mutations by pinpointing whether they are located in exons vulnerable to ESMs.

## Introduction

With the increasing use of genome wide sequencing, detection of variants is now widely implemented in routine diagnostics (1). Whereas interpretation of the effect of classical missense and nonsense mutations that directly affect the amino acid sequence of a protein appears straightforward, identification and characterization of exonic mutations that alter the splicing code (2,3) is still challenging.

Correct assessment of the impact that a mutation may have on the splicing code is vital for correct classification of variants and numerous *in silico* tools have been developed to help predicting this (4-7). These tools have been widely implemented diagnostically to enable clinicians to correctly call the pathogenicity of identified sequence variants and make more accurate decisions (8). Mutations located in the splice site regions at the terminal parts of exons are now often correctly recognized as splice altering rather than amino acid altering (9), but variants located outside of the canonical splice sites are often misclassified when based solely on their effect on the amino acid code (10). Several tools have been developed for analysis of a mutation’s effect on the splicing regulatory elements (SRE) (11). These tools estimate the impact of a mutation, either through an analysis of known SRE sequences, or more recently, by incorporating machine learning to account for the large variety of features that may be impacted by the mutation (5,12,13). There are many challenges to accurately predict splicing mutations using such *in silico* tools, and often the predictions do not accurately or consistently predict the effect of a mutation so that this mimics the observed *in vivo* effect on splicing (14). One of the primary shortcomings of the current *in silico* tools is the lack of consideration of genomic context dependent effects and the complex interplay between genomic context, splice site strengths and SREs, which is influenced by numerous parameters. Although empirical data from patient cells naturally include all information and therefore is superior to models, *in vivo* validation of splicing is often limited by availability of patient tissue samples or requires generation of a suitable splicing reporter able to mimic the endogenous context accurately.

Importantly, it is usually assumed that all exons have an equal dependency on SREs and consequently that sequence variants that alter SREs are equally likely to cause aberrant splicing in all exons. Despite this, studies employing model exons demonstrate that some exons can be skipped by mutations located across the entire exon, while others are only affected by mutations located in close proximity to the splice sites (15). Further, it has recently been reported that up to 77% of exonic mutations in *MLH1* exon 10 and 60% of exonic mutations in *BRCA2* exon 3 affect splicing (9,10), suggesting that the majority of exonic mutations affect splicing and that this is a very frequent disease mechanism. In sharp contrast to this, another study reported that only 10% of 4,964 exonic disease associated mutations altered splicing (16). It is at present unknown what causes these large differences in the reported proportion of exonic mutations that affect splicing of an exon and whether 10% or 70% should be expected when these findings are extrapolated to other exons. We have previously demonstrated that *ACADM* exon 5 is particularly vulnerable to splicing mutations and that identical mutations in SREs have different effects dependent on the exon where they reside, as well as the genomic context (17). We hypothesize that such different vulnerability of exons may explain the widely different proportions of exonic splicing mutations (ESMs) observed in the studies above. Based on our observations from *ACADM*, and other genes harboring particularly vulnerable exons, we speculated that this vulnerability can be revealed as a low degree of exon skipping from normal cells. Therefore, we developed VulExMap that designates whether an exon is vulnerable, based on junction counts from Snaptron GTEx data (18). We used VulExMap to show that, on a transcriptome-wide basis, vulnerability to ESMs differs between constitutive exons. Importantly, we collected 331 previously reported ESMs from 71 genes and demonstrate that ESMs are mainly located in the small proportion of exons predicted to be vulnerable by VulExMap.

## Materials and methods

### Discovery of vulnerable exons in RNA sequencing data – VulExMap

The VulExMap tool is divided into tabs, which each is a layer deeper into the process of analyzing the data. The first tab is for uploading data (for now it is only possible to upload a BED file containing SNPs, all other files are preloaded files used when analyzing the data for this paper). The second tab displays all possible genes and let you choose one to plot, and the third is a graph of the gene showing all the exons and color coding them according to vulnerability. To identify vulnerable exons, we downloaded junction counts from the Snaptron server (18) belonging to the GTEx samples (19), and estimated inclusion levels using flattened gene models of the hg38 NCBI refGene annotation downloaded from UCSC (20). SAJR (21) was used to generate the gene models and using its splice-site annotation of gene segments we computed inclusion estimates as outlined in Supplementary figure S1. To eliminate low coverage samples, only samples where segments with ≥ 10 combined inclusion and exclusion junction counts were included in the final estimates of inclusion of each segment, and only cassette exons with a mean junction read count of at least 30 were used in downstream analyses. We then calculated the coefficient of variation (CV) of the PSI for each cassette exon as the standard error of the mean divided by the mean of the PSI across the samples. Next, we divided exons into bins based on the mean PSI so that we obtained bins of cassette exons with PSI in the ranges [0-80[, [80-85[, [85-90[, [90-95[, [95-96[, [96-97[, [97-98[, [98-99[, [99-99.5[, and [99.5-100]. We then defined thresholds of minimum and maximum CV for vulnerable cassette exon using the median of the CV of cassette exons in the [85-90[PSI bin as the maximum and the median of the CV in the [99-99.5[PSI bin as the minimum. We then defined resilient exons to be exons that were robustly included in all sequenced samples, i.e. PSI (Percent spliced in) > 99% and CV < minimum CV threshold, while exons that had a higher deviation of inclusion between samples resulting in a CV value between the two CV thresholds and a PSI > 85%, were defined as vulnerable. All other cassette exons were categorized as alternative, unless their PSI was exactly 0, in which case we classed them as not spliced in. Exons with junction counts below the threshold was classified as NA. For the online webservice we applied these classification criteria to all segments, including segments that are not cassette exons, but in the analyses in the manuscript we exclusively considered cassette exons as defined in the refGene annotation. The online VulExMap webtool is available at https://vulexmap.compbio.sdu.dk.

### Sequence analysis of resilient and vulnerable exons

The vulnerable, resilient, and alternative segments for each dataset was compared by characteristics associated with exon definition. We used PESE and PESS motif databases (22) to identify general ESE and ESS motif density. We used MaxEntScan (23) to obtain maximum entropy scores of the donor and acceptor sites as a measure of splice site strength. Up- and downstream splice sites were obtained using the immediate up- and downstream exons annotated in the hg38 refGene annotation table from UCSC (20). K-mer analysis was performed with jellyfish 2.2.6 (24). Multiple transcripts in some genes were resolved by using the transcript with the lowest identification number.

### ESM Database

To collect all previously reported ESMs, pubmed was searched using the search terms “exonic splicing mutation” and “exon skipping minigene assay”, as well as inhouse knowledge of ESM reports. All reported ESMs was then manually curated in ENSEMBL to the canonical refseq transcript. Mutations located in the first or last three bases of an exon were excluded from the database, as these were presumed to disrupt the splice site sequences. Furthermore, mutations that were reported to create a new splice site were also excluded. Finally, mutations in annotated alternative exons were also excluded from the database. All mutations in table S1 were, if necessary, reassigned correct HGVS nomenclature and position on hg38 is included. The database was last updated on November 10^th^ 2022.

### Minigenes

*ACADM* exon 2, *BRCA2* exon 3 and *ATM* exon 40 was cloned into the pSPL3 splicing reporter, a vector developed for exon trapping (25). Inserts was amplified from genomic DNA using primers in the introns flanking the selected exons; MCEX2S: Xhol: 5‘-CTGTACAAGGACTCGAGATAACTGATAATTGGCT-3’, MCEX2AS-BamHI: 5‘-GGACAGTGGATCCATTCTACTCATTGAAAGACA-3’, BRCA2_ex3_XhoI_XbaI_F: TACGACTCGAGTCT-AGATGGCCGAATTTTATCGTGGAA, BRCA2_ex3_EcoRI_BamHI_R: TACGAGGATCCGAATTCGCACCTACGCCA-GGGAAA, ATM_ex40_ XhoI_XbaI_F: TACGACTCGAGTCTAGATGAATTGGATGGCATCTGCTCT and ATM_ex40_ EcoRI_BamHI_R: TACGAGGATCCGAATTCGGCAAGCATCCCAGACAGTA. *BRCA2* exon 3 and *ATM* exon 40 cloning primers were designed with an additional restriction site, for late subcloning into the *ACADM* minigene (17). Cloning into the pSPL3 vector was confirmed with Sanger sequencing (Eurofins) and mutagenesis was performed by Synbio technologies. After mutagenesis, the *BRCA2* exon 3 and *ATM* exon 40 inserts were subcloned into the *ACADM* minigene with either the normal or the optimized downstream 3’ss, between XbaI and EcoRI restriction sites. Subcloning was performed by Synbio technologies.

### Transfection of cells

HeLa cells were cultured in RPMI 1640 medium (Lonza, Copenhagen, DK) supplemented with 10% (v/v) fetal calf serum, 0.29 mg/ml glutamine, 100 U/ml penicillin, and 0,1 mg/ml streptomyocin at 37°C in 5% (v/v) carbon dioxide. The cells were grown to ∼75% confluence in 12 well plates and transfected with 0.2 μg minigene plasmid. Transfections were carried out in two biological replicates with duplicates.

### RNA extraction and cDNA synthesis

After 48 hours the cells were lysed using 0.5mL Trizol® reagent (Fisher) pr. well. Total RNA was extracted using chloroform and precipitated with isopropanol. Complementary DNA (cDNA) was synthesized from 0.5 μg RNA using the High capacity cDNA kit (Invitrogen).

### PCR analysis of splicing

To investigate the mutations effect on splicing, plasmid specific primers were used. In pSPL3 we used the primers SD6: 5’ TCTGAGTCACCTGGACAACC and SA2: 5’ ATCTCAGTGGTATTTGTGAGC. In the *ACADM_*pcDNA minigenes the primers were located in; exon 4 forward primer: 5’CCTGGAACTTGGTTTAATG and exon 6-pcDNA reverse primer: 5’ AGACTCGAGTTACTAATTAATTACACATC. The cDNA was amplified using TEMPase HOT Start DNA polymerase (Ampliqon). The PCR program consisted of 15 min at 95°C, followed by 32 cycles of 30 sec at 95°C, 30 sec at 55°C and 30 sec at 72°C, followed by 5 min at 72°C. After PCR, the samples were visualized by agarose gel electrophoresis using SeaKem LE agarose (Lonza) and Gelred (Biotium). Exon skipping was quantified using the DNF-910 dsDNA analysis kit (35-1500 bp) on a Fragment analyzer (Advanced Analytical).

### Surface plasmon resonance imaging

Surface plasmon resonance imaging was carried out as previously described (26). Briefly, biotinylated RNA-oligonucleotides were immobilized onto a G-strep sensor chip (SSENS) for 20 min. The following recombinant proteins was injected for 8 minutes, followed by dissociation for 4 minutes; SRSF1 (Genscript) and hnRNPA1 (Abcam, ab123212). Nuclear extract was used as a control of the oligoes binding efficiency. Binding was fitted to a 1:1 kinetics model with Scrubber2 (v. 2.1.1.0; Biologics inc.). For hnRNPA1 a biphasic 1:2 model was used in ClampXP (version 3.50; Biosensor Data Analysis).

### RNA affinity purification

Affinity purification of RNA-binding proteins was performed as previously described (17) using biotinylated RNA oligonucleotides (LGC biosearch technologies, Risskov, Denmark, sequences in supplements) immobilized on Dynabeads M-280 streptavidin magnetic beads (Invitrogen) and incubated with HeLa cell nuclear extract. The proteins was separated on a precast 4-12% NuPAGE Bis-Tris gel (Invitrogen) and transferred to an Immobilion-PDVF membrane (Millipore). The membrane was then blotted with antibodies against SRSF1 (Zymed/Invitrogen, 535814A), and hnRNPA1 (Sigma-Aldrich, 080M4857), and hnRNPL (Santa Cruz, sc-32317) as a binding control. Secondary antibodies were either Goat-Anti-Mouse (ThermoFischer Scientific, A16066) or Goat-Anti-Rabbit (ThermoFischer Scientific, A16104).

## Results

### Vulnerable exons can be detected empirically with VulExMap

We previously defined *ACADM* exon 5 as a vulnerable exon, because disruption of any of several SREs by disease associated exonic point mutations can affect exon inclusion (17). Interestingly, it has previously been observed that *ACADM* exon 5 is skipped in a small proportion of cDNA from patient and control cells (27,28). We noted that low levels of background skipping have also been reported in other genes, especially in exons with a high occurrence of ESMs (29-32). We therefore hypothesized that low levels of skipping could be a general indicator of exon vulnerability to ESMs. To test this hypothesis, we developed VulExMap. This tool can identify vulnerable exons based on analysis of RNA-seq junction count data, allowing the creation of a map of vulnerable exons on a genomic scale, to help researchers evaluate if mutations that affect SREs are likely to cause exon skipping. In order to establish VulExMap we used Snaptron GTEx junctions (18), because a large sample size is necessary in order to enable detection of the small, but significant levels of exon skipping that indicate exon vulnerability. Based on inclusion levels (PSI) and coefficient of variation (CV), VulExMap categorizes exons as either vulnerable, resilient, or alternative (figure 1A). To ensure robustness we only include exon segments with a mean combined inclusion and exclusion junction count ≥ 30, and a minimum junction read count of 10 reads in VulExMap. We developed an online web-tool available at https://vulexmap.compbio.sdu.dk, where VulExMap can be used to analyze any gene of interest to detect vulnerable exons (supplementary figure S2).

**Figure 1.**
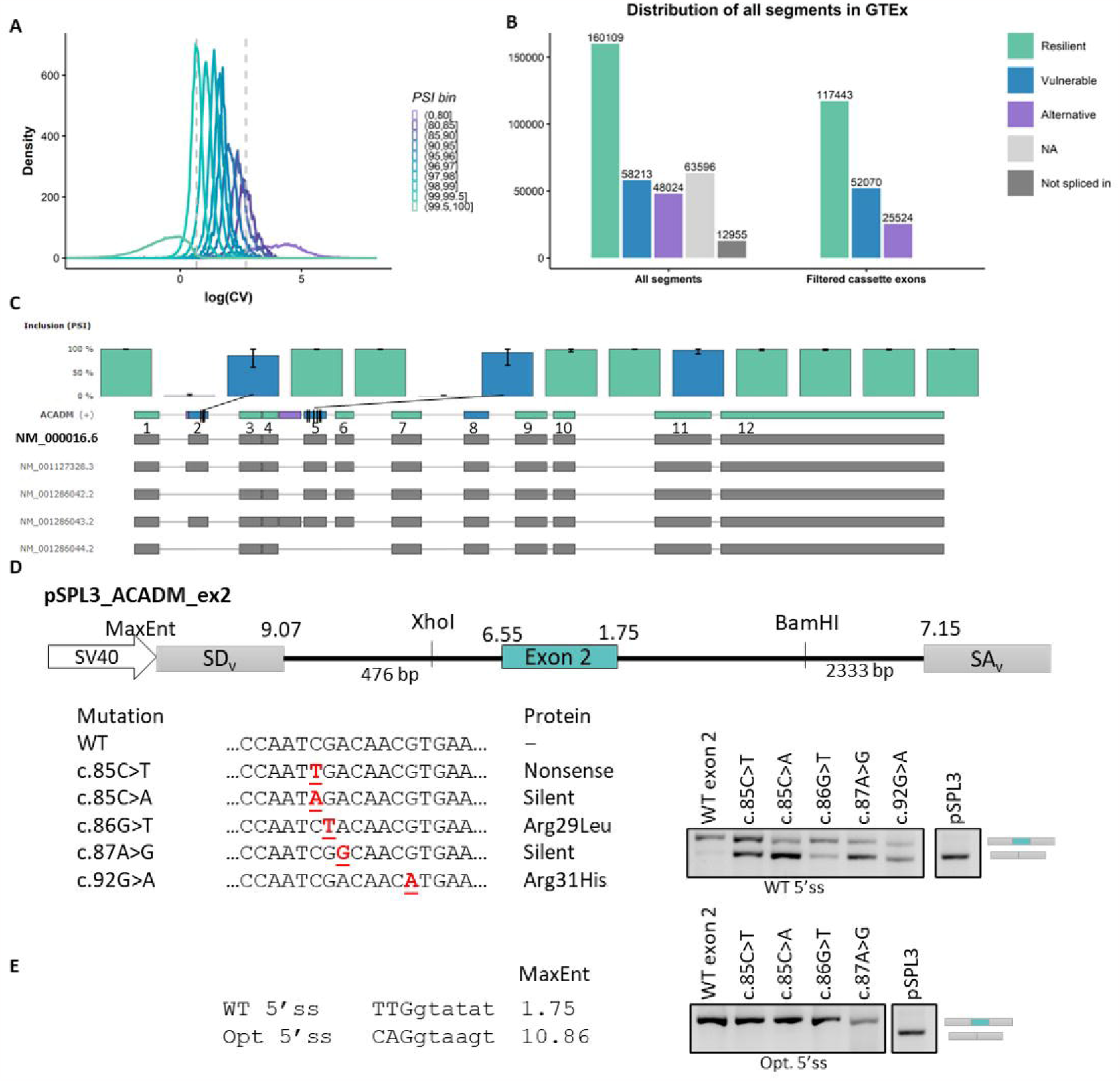
VulExMap reveals vulnerable constitutive exons. **A**. Logarithm of the coefficient of variation (CV) of binned internal exons with sufficient reads. Exons were binned into groups based on mean PSI level and CV thresholds were established as the median CV value of the exons with PSI in the ranges of 80-85% and 99-99.5% (indicated in dashed vertical lines). **B**. Distribution of segments from GTEx classified by VulExMap. The segments were classified based on PSI and CV, and only included if the segments had more than 50 junctions covering the exon in more than 30 samples. Exons with a PSI > 99.5 were classified as resilient, exons with a PSI <99.5 and >85 were classified as vulnerable, and exons with a PSI <85 were classified as alternative. If an exon had a PSI = 0, it was not spliced in, and if the exon was not extensively included, if was termed NA. **C**. VulExMap of *ACADM* reveal 3 vulnerable exons. The primary protein coding refseq transcript NM_000016.1 is shown directly below the gene model. **D**. pSPL3 splicing reporter with the vulnerable *ACADM* exon 2 was used to investigate mutations from patients with MCAD deficiency. RT-PCR was carried out on cDNA from HeLa cells transfected with the plasmids. **E**. Optimized 5’ss was introduced to some of the vectors which was transfected as above.

With these threshold settings we find that on a global scale, 27% (52,070) of exons with junction data in Snaptron (18) are vulnerable to ESMs (figure 1B). When we used VulExMap to analyze *ACADM* (figure 1C) this supported that *ACADM* exon 5 is in fact vulnerable. Interestingly, VulExMap also categorizes *ACADM* exon 2 as vulnerable, and a silent c.85C>A mutation in this exon has been reported to cause disease (33-36). To investigate this, we therefore cloned *ACADM* exon 2 and flanking intronic sequences into a splicing reporter vector with either the wild type (WT) sequence or containing mutations identified by newborn screening to cause MCADD (33) (figure 1D). Consistent with the predicted vulnerability, we observed exon 2 skipping from several mutations in this exon, including the two c.85C>A and c.87A>G silent mutations. Analysis of patient lymphoblasts from a patient harboring the c.87A>G mutation as one of the disease alleles showed that this mutation does indeed cause exon 2 skipping also in the endogenous gene (Supplementary figure S3). As the 5’ss of exon 2 is weak (MaxEnt 1.75), we strengthened the score of the 5’ss (MaxEnt 10.86) in the splicing reporter to see if this would make exon 2 resilient and thereby less responsive to the mutations. When optimizing the 5’ss we observed complete exon inclusion from all mutations (figure 1E). Interestingly, mutations altering the motif ATCGACA (*ACADM* cDNA pos 83-89) also result in exon skipping in other genes, suggesting that this motif functions as an ESE. In *F9* exon 5 mutation of the C corresponding to pos. 85 to all three different possibilities and the C corresponding to pos. 88 to T has been reported to cause exon skipping, with the C>A mutation causing the most dramatic effect (37). This is consistent with our observation from *ACADM* exon 2, where c.85C>A causes a more severe exon skipping than c.85C>T (Figure 1D) and might suggest that the C>A mutation causes a stronger effect by simultaneously disrupting an ESE and creating an ESS, whereas the other variants only disrupt the ESE. In support for this, the c.840C>T mutation in *SMN2* creates exactly the same TAGACA hnRNPA1-binding ESS motif (26,38-40) that is created by the C>A mutation in *ACADM* exon 2 and *F9* exon 5, showing that these mutations have a dual effect by simultaneously abolishing an ESE and creating an ESS.

### Vulnerable exons share multiple characteristics with alternative exons

Because our previous experimental analysis of *ACADM* (17) showed that vulnerability is determined by several factors, like SREs, splice site strength and the genomic context, we compared exons which are classified as either vulnerable (n = 52,315), resilient (n = 109,460), or alternative (n = 26,091). We first scored the SRE density in the three groups. We used the sample of 2,069 putative exonic splicing enhancers (PESE) and 974 putative exonic splicing silencers (PESS) from Zhang et al. 2004 (41) (figure 2A-B). Interestingly, this showed that there are significantly fewer ESEs/bp in the vulnerable exons compared to the resilient exons (p < 2.22e-16, Wilcoxon rank sum). Conversely, the vulnerable exons have a significantly higher ESS density than the resilient exons (p < 2.22e-16). Analysis of GC content revealed that vulnerable exons also have a significantly lower GC content (p < 0.0005) than resilient exons, which further supports the notion that exon definition is weaker for vulnerable exons (figure 2C), since GC content is directly linked with exon definition (42). The vulnerable exons were also shorter than the resilient exons (figure 2D) and were flanked by longer introns.

**Figure 2.**
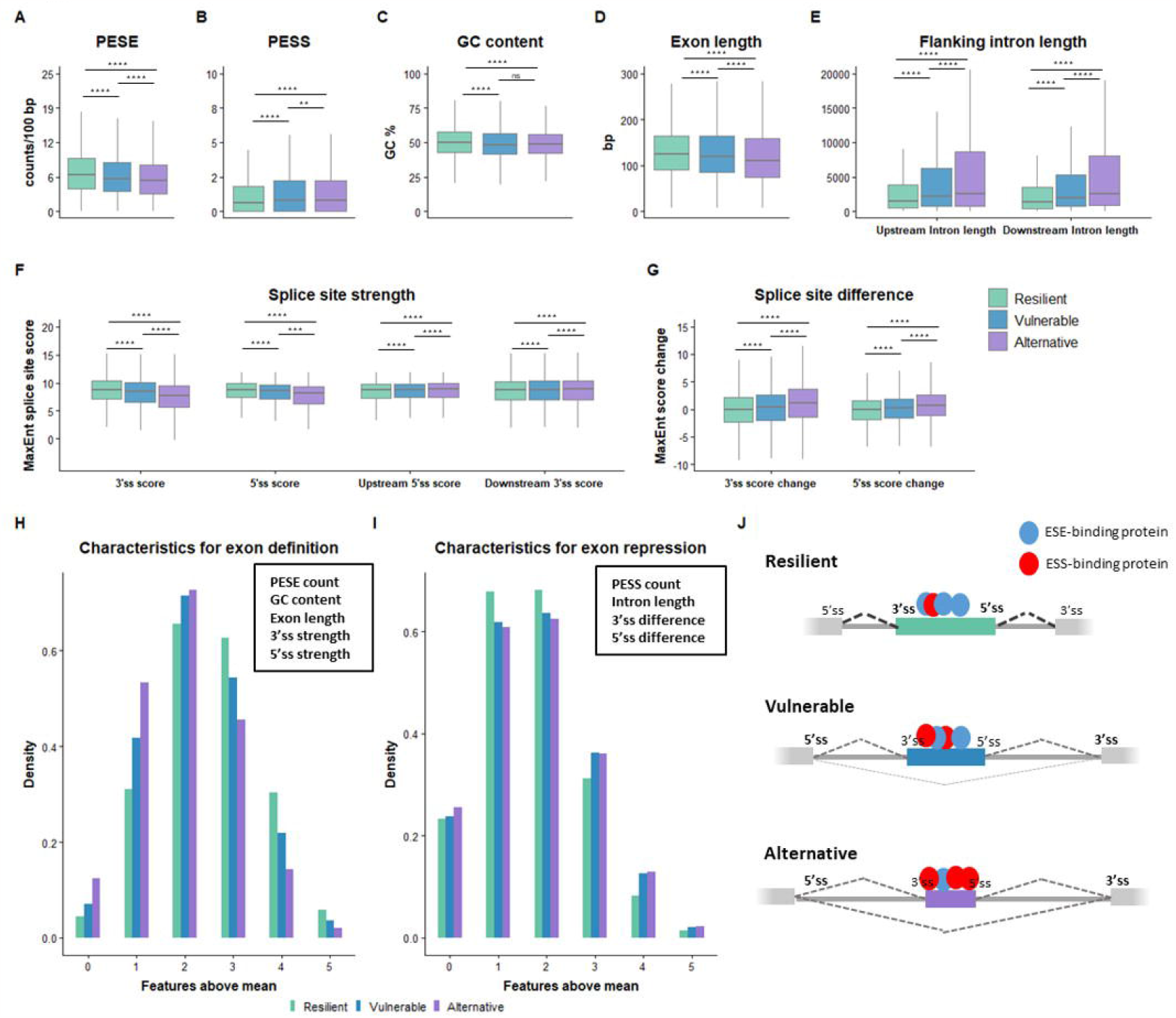
From the GTEx RNA-seq data segments that were identified as vulnerable (vulnerable (n = 52,315), resilient (n = 109,460), or alternative (n = 26,091) was included for comparison. Only internal exons were included. **A-G**. In each group PESEs/100 bp (**A**) and PESSs/100 bp (**B**) and GC content (**C**) was calculated. To determine context, we also compared exon length (**D**), flanking intron length (**F**) and splice site strength (**F**). To acquire the difference between to the flanking up – and downstream splice sites, we took the 3’ss score of the downstream 3’ss and subtracted the 3’ss score of the exons in each group, or the 5’ss score of the upstream 5’ss and subtracted the 5’ss score (**G**). Wilcoxon rank sum test was performed between the three groups. **** p ≤ 0.00005, *** p ≤ 0.0005, ** p ≤ 0.005, ns p > 0.05. F. **H - I**. Density of features associated with exon definition (**H**) of exon repression (**I**). The mean score of each feature above was calculated, and the exons with a score above the total mean was grouped by whether they had 0, 1, 2, 3, 4 or 5 different feature scores above the total mean. **J**. Graphical representation of the many factors that can affect exon definition. Resilient exons are, in general, long, with a high density of ESEs and low density of ESSs, strong splice sites compared to the flanking exons’ splice sites, and shorter introns. Vulnerable exons are weaker on multiple parameters than resilient exons, but more defined than alternative exons.

Analysis of splice site strength shows that the vulnerable exons have significantly weaker 3’ss and 5’ss than the resilient exons (figure 2F). This is very interesting, as it has previously been reported that exons with weak splice sites generally have a higher density of ESEs in order to achieve sufficiently strong exon definition (43), whereas we observe the opposite for the group of vulnerable exons, indicating that their vulnerability may in part be caused by a low density of ESEs, which is very close to be insufficient to compensate for the weaker splice sites. Furthermore, the proximal splice site of the exons that flank the vulnerable exons are stronger than the splice sites of the vulnerable exons, indicating that competition with the downstream and upstream splice sites are also important in defining vulnerability of an exon (figure 2G).

Because vulnerability of each individual exon is defined by several features each contributing with different weight to the overall vulnerability of that individual exon there are exons in each group (vulnerable, resilient and alternative) with scores above or below the mean for a feature in another group. Therefore, we separated the different features into positive (PESE, GC content, exon length, splice site strength) and negative (PESS, flanking intron length, difference from flanking exon splice site) characteristics associated with either exon inclusion or splicing repression. We then took the total mean for all exons in all groups and counted the density of exons in each group with a score above the mean of a certain feature. When grouping the density of the features, we observe, that for the positive characteristics, the alternative exons had the highest density for few characteristics above mean, whereas the resilient exons had the highest density for having more than two characteristics associated with exon inclusion (figure 2H). Conversely, the resilient exons had the highest density for having 0, 1 or 2 characteristics associated with repression, whereas the vulnerable and alternative exons both had the highest density of negative features (figure 2I). This indicates that resilient exons are innately stronger defined than vulnerable exons, but they may still contain few features normally associated with weak exons. Similarly vulnerable exons have fewer positive characteristics than resilient exons, but are more strongly defined than alternative exons (figure 2J).

### Exonic splicing mutations are overrepresented in vulnerable exons

Based on our findings we speculated whether vulnerable exons are more sensitive to changes to the splicing code caused by ESMs. To investigate if mutations in vulnerable exons are more likely to cause exon skipping, we first manually searched the literature for ESMs that had been functionally validated to cause exon skipping (figure 3A). Exonic mutations in the splice sites, i.e. the first three and last three nucleotides of the exon, were excluded from the analysis, as these would typically result in exon skipping regardless of the vulnerability of the exon. Mutations that activate cryptic splice sites, as well as mutations in alternative exons, were also excluded, as we only wanted to establish the distribution of mutations affecting constitutively spliced exons. In total, 331 ESMs were included in our database, of which 313 are single nucleotide variants (supplementary table S1). Next, we analyzed the distribution of these ESMs in the exons classified by VulExMap. Interestingly, 236 of 330 ESMs (72%) were located in vulnerable exons, whereas only 94 of 330 ESMs (28%) were located in resilient exons (figure 3B). Only one ESM could not be classified, due to the lack of *USH2A* expression in GTEx. In total there were 63 (65%) different vulnerable exons and only 33 (35%) resilient exons in the ESM database (figure 3C), although vulnerable exons only make up 27% of all exons (19% in the genes in the database, figure 3D). There is a statistically significant difference (p = 3.37e-23) in the proportion of vulnerable exons between exons harboring ESMs (63/96 exons were vulnerable) and vulnerable exons in general within the same genes harboring ESMs (843/4425 exons were vulnerable). With a 3.41-fold relative difference and an odds ratio of 4.87 [3.47; 6.83] this suggests that exonic mutations are three times more likely to affect splicing when located in a vulnerable exon than when located in a resilient exon. Due to this distribution, we wanted to know if vulnerable exons in general contained more mutations, and whether these were associated with aberrant splicing. When mapping the GTEx sQTLs (19) and dnSNP156 variants (44) to exons classified by VulExMap, we observed that vulnerable exons had twice as many sQTLs/SNP as resilient exons (figure 3E, supplementary figure S4). Alternative exons had the highest number of both sQTLs and SNPs in general, which indicates a higher tolerance towards missense variants. Furthermore, vulnerable exons have slightly fewer synonymous variants than resilient exons (supplementary figure S3), which may represent an evolutionary constraint against mutations, as silent mutations have a higher chance of causing exon skipping in vulnerable exons. We have linked the ESM database with VulExMap, so that all ESMs from the database are automatically displayed in the corresponding exons at https://vulexmap.compbio.sdu.dk.

**Figure 3.**
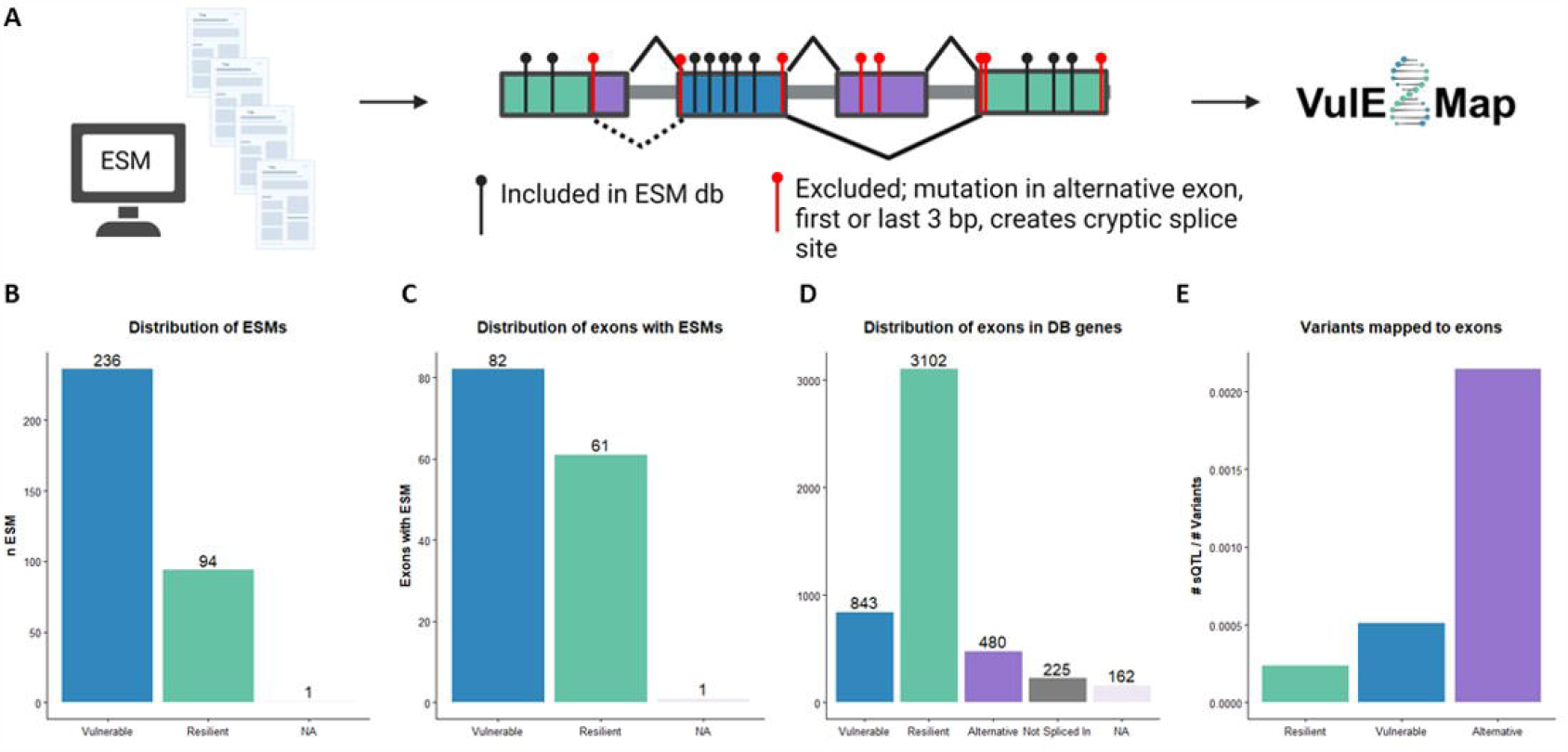
Vulnerable exons are enriched with ESMs and predisposed to skipping. **A**. The ESM database was generated by manually searching PubMed using the search terms “exonic splicing mutation” and “exon skipping minigene assay”. Mutations were assigned correct HGMD nomenclature. Mutations in alternative exons, the first and last 3 bp of an exon or mutations that created new splice sites, were excluded from the database. All mutations in the ESM database were uploaded to VulExMap. Figure made with BioRender. **B**. Distribution of 331 ESMs in vulnerable and resilient exons, from 72 genes. VulExMap was used to classify the exons containing the ESMs from table S1. If the gene had a mean segment count < 30 in all five datasets, then those ESM could not be classified = NA (only *USH2A* could not be classified due to low expression in GTEx). **C**. Classification of exons with ESMs, eliminating any bias for multiple ESMs in the same exon, still show an overrepresentation of ESMs in vulnerable exons. **D**. Distribution of all exons in the 72 genes from the ESM database that could be classified according to the predetermined threshold values. **E**. Distribution of all dbSNP156 variants, outside of the first and last 3 bases of the exon, in GTEx sQTLs show that vulnerable exons have more SNPs associated with splicing.

### Exonic splicing mutations affect similar motifs in vulnerable exons

Although there are common motifs for the splicing regulatory elements in different exons, the consequences on splicing from abrogating identical SREs in different exons is not necessarily the same (17). To identify essential elements for exon definition, we performed K-mer analysis of enriched hexamers in the vulnerable and resilient group of exons. Interestingly, the most enriched hexamer in the resilient exons was the GAAGAA ESE (figure 4A), which we have previously demonstrated to be critical for inclusion of the vulnerable *ACADM* exon 5 (17). Furthermore, we observe that the top hexamers in the resilient exons were all highly represented in the PESE database (supplementary table S2). When comparing to the alternative exons, a wider dispersion of K-mer z-scores was observed (Supplementary figure S4). Although the K-mer analysis revealed that the GAAGAA ESE is the most enriched hexamer in the resilient exons, we observed that the ESM database (supplementary table S1) contains 10 mutations that disrupt the GAAGAA motif located in vulnerable exons. The only example of a resilient exon being skipped by a mutation disrupting a GAAGAA motif is *ATM* exon 40 (NM_000051:c.5932G>T). The GAAGAA hexamer has previously been identified as an SRSF1-binding ESE (45), which is consistent with our previous work with the GAAGAA ESE in *ACADM* exon 5 (17). None of the ESMs created the GAAGAA motif. Furthermore, GCTGGG and CTGGGG hexamers were enriched in the vulnerable exons (figure 4A). The CTGGGG motif was created by three different ESMs whereas the GCTGGG was created by three ESMs and abolished by two ESMs. Most likely, these two hexamers are enriched because they harbor GGG triplet motifs, which are known to function as ESS’s (46,47). In fact, 27 (8%) of the ESMs created triplet GGG motifs, whereas only three ESMs disrupted triple GGG motifs. In two instances, we suspect that a cryptic donor splice site is generated as GG from the GGG triplet changes to GT. In the last example a TGGG is altered, so that it instead constitutes the stronger TAGG ESS motif, which is a well-established hnRNPA1 binding motif (40). In total, 17 ESMs resulted in creation of the TAGG ESS motif, whereas this motif was not disrupted by any ESM. To investigate if indeed all GAAGAA-disrupting mutations in the ESM database affect SRSF1 binding and that the mutations creating TAGG all affect hnRNPA1 binding we performed SPRi analysis employing biotinylated RNA-oligonucleotides, with either WT or mutant 15-mer sequences (figure 4B, supplementary table S3). For SRSF1 we observe a significant reduction of binding to the GAAGAA oligonucleotides, when the GAAGAA motif is disrupted (P = 0.027, Wilcoxon signed rank). The only case where the mutation actually increases binding of SRSF1 is *MFSD8* c.750A>G, where the mutation changes GAAGAA to GAGGAA, which is still considered a strong SRSF1-binding ESE according to DeepCLIP analysis (48) (supplementary table S4). Our analysis showed that the exon skipping effect of this mutation is instead caused by a simultaneously increased binding of hnRNPA1 to an already strong ESS (figure 4B, supplementary table S3 and S4). In the only resilient exon with a GAAGAA-disrupting mutation, *ATM* exon 40 (NM_000051:c.5932G>T), we observe simultaneous disruption of SRSF1 binding and a simultaneous increase in hnRNPA1 binding consistent with a dual effect from loss of an ESE and gain of an ESS (figure 4B, supplementary table S3). This indicates that the resilient *ATM* exon 40, requires a larger, dual change in exon definition to be skipped.

**Figure 4.**
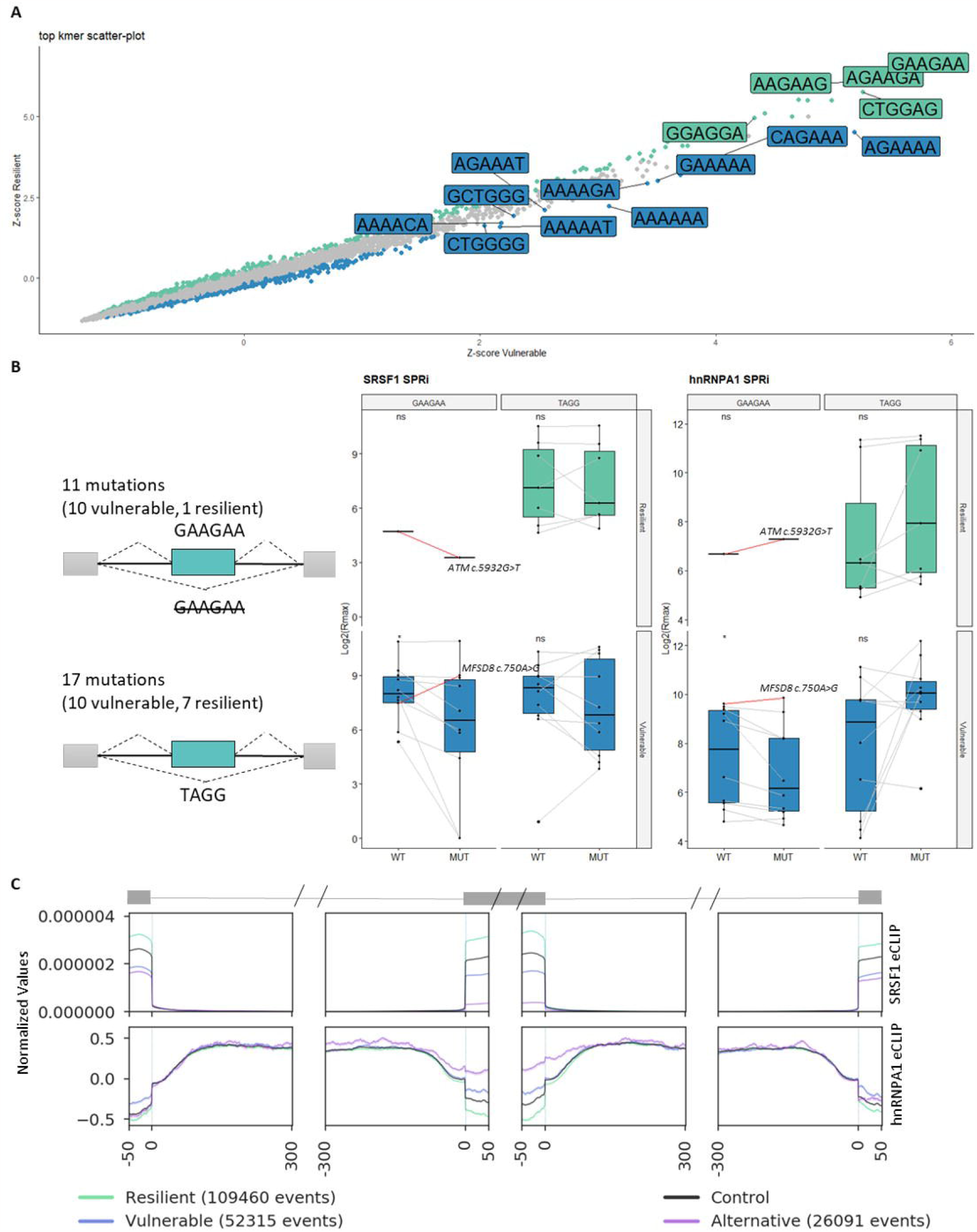
Overlapping motifs indicate exon vulnerability. **A**. K-mer analysis (jellyfish) was carried out on the sequences of the exons that were either vulnerable or resilient, and differentially enriched hexamers were compared between the two groups. Shown are the top enriched hexamers in the vulnerable (blue) and resilient (green) exons. Significantly enriched k-mers were identified using a proportion test and adjusting for multiple testing with Bonferroni. **B**. 11 mutations affecting the GAAGAA motif, and 17 mutations creating a TAGG ESS motif was introduced in 15-mer biotinylated RNA oligonucleotides, and the wild type (WT) and mutant (MUT) sequences was analyzed with surface plasmon resonance imaging (SPRi) using recombinant SRSF1 and hnRNPA1. Binding is shown as Log2(Rmax). Red lines indicate the two mutations in *ATM* and *MFSD8*. Wilcoxon paired test was performed between WT and MUT binding scores. * p ≤ 0.05, ns p > 0.05. **C**. ENCODE eCLIP data from K562 cells against SRSF1 and hnRNPA1 was plotted across the last 50 bp of the upstream exon, the first and last 300 bp of the upstream intron, the first and last 50 bp of the resilient vulnerable and control exons, the first and last 300 bp of the downstream intron, and the first 50 bp of the downstream exon. Control is the mean binding of all exons in the three groups.

The TAGG-creating mutations all increase binding of hnRNPA1 in the resilient exons, with the exception of *PYGM* c.1085G>A, where the TGGG to TAGG instead increases binding of hnRNPA2 (Rmax increased from 135.4 to 186.9, supplementary table S3). Furthermore, the TAGG mutations created in the vulnerable exons do not all increase binding of hnRNPA1, but some instead increased binding of hnRNPA2 or hnRNPH (supplementary table S3). The overlapping ESS motifs increases the likelihood of a mutation creating an ESS, while simultaneously disrupting an ESE.

Interestingly, the SPRi analysis showed that the GAAGAA mutations in the vulnerable exons reduced binding of hnRNPA1, despite the mutations causing exon skipping (figure 4B). This illustrates how the balance of splicing factors is not 1:1 but loss of both a negative and positive element can still result in aberrant splicing in vulnerable exons.

To validate the importance of SRSF1 and hnRNPA1 as splicing regulators in the endogenous context, we mapped ENCODE eCLIP data (49) from K562 cells across all the exons classified by VulExMap (figure 4C). Consistent with the PESE and PESS distribution (figure 2A+B), we observed higher distribution of SRSF1 eCLIP reads across the resilient exons, and the highest distribution of hnRNPA1 eCLIP reads across the alternative exons, with the vulnerable exons having less hnRNPA1 eCLIP reads than alternative exons, but more hnRNPA1 eCLIP reads than both alternative and resilient exons in the flanking up- and downstream exons, underscoring the importance of context on exon vulnerability.

These data indicate that vulnerable exons are defined by a very finely tuned balance between positive and negative SREs, which can be easily disrupted by a point mutation that either abolishes an ESE or creates an ESS, whereas the balance in resilient exons is less fragile and require stronger shifts, possibly due to the context in which the exon gains its resilience to aberrant splicing.

### Vulnerability to splicing mutations is context-dependent

We have previously shown that the vulnerable *ACADM* exon 5 is strongly affected when inserted into a different context, whereas the resilient *ACADM* exon 6 was not affected when inserted in a similar context (17). Due to the high frequency of mutated GAAGAA motifs in the ESM database, we chose to investigate the vulnerable *BRCA2* exon 3 (NM_000059) and the resilient *ATM* exon 40 (NM_000051), which are both skipped by mutations disrupting a GAAGAA ESE. We chose two mutations from *BRCA2* exon 3, namely the c.100G>A mutation which disrupts SRSF1 and hnRNP A1 binding completely in the SPRi experiments (supplementary table S3), and the c.145G>T mutation, which simultaneously disrupt SRSF1 binding and only partially decreases hnRNPA1 binding to a strong ESS in the SPRi data and increases hnRNPA1 according to DeepCLIP analysis (supplementary table S4, supplementary figure S6). In the resilient *ATM* exon 40, only the c.5932G>T mutation has been reported to cause exon skipping by disrupting a GAAGAA ESE, which we demonstrate both disrupt SRSF1 binding and create hnRNPA1 binding with SPRi (figure 4B, supplementary table S3) We therefore designed an artificial c.5935G>A mutation (GAAGAA to GAAAAAA), which is predicted to not only affect SRSF1 binding, but also decrease hnRNPA1 binding using DeepCLIP (supplementary tableS4, supplementary figure S7). We used RNA affinity pulldown analysis to confirm these effects of the two *BRCA2* (figure 5A) and *ATM* (figure 5B) mutations, and the effects were also consistent with the results from SPRi analysis. Therefore we hypothesize that *BRCA2* c.145G>T and *ATM* c.5932G>T will have a more severe effect on splicing, due to the simultaneous ESE loss + ESS gain dual effect.

**Figure 5.**
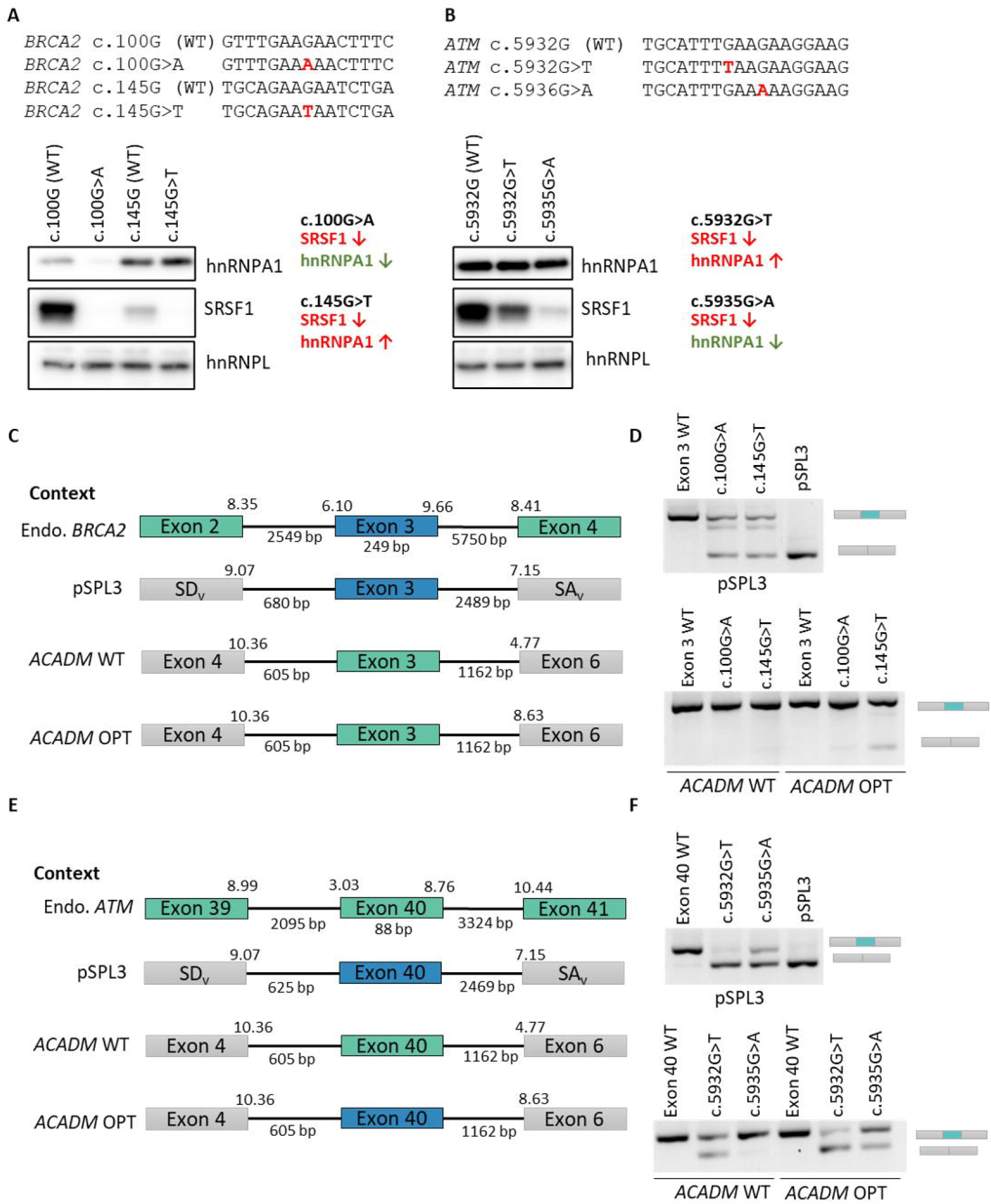
Exonic context determines splicing outcome in vectors. **A**. Two mutations in the vulnerable *BRCA2* exon 3 were selected and the effect the mutations had on protein binding was verified with RNA-affinity pulldown with nuclear extract. hnRNPL was included as a binding control. The c.100A>G mutation is considered a mild mutation, and the c.145G>T mutation is severe. **B**. In the resilient *ATM* exon 40 there has only been reported one splicing mutation, c.5932G>T. a second mutation at c.5935G>A was included, as this was predicted to be less severe than the c.5932G>T (supplementary figure S5). Pulldown with nuclear extract confirmed that c.5932G>T was severe and c.5935G>A was mild. **C**. The vulnerable *BRCA2* exon 3 with WT, c.100G>A or c.145G>T was introduced into three different vectors, the pSPL3 splicing reporter or the *ACADM* exon 5 minigene from (17) with either WT or optimized (OPT) downstream 3’ss. Shown is also the context in endogenous *BRCA2*. **D**. The splicing reporters with *BRCA2* exon 3 were transfected into HeLa cells, and splicing was evaluated with RT-PCR. **E**. The Resilient *ATM* exon 40 with WT, c.5932G>T or c.5935G>A was introduced into the same splicing reporters as *BRCA2* exon 3. Shown is also the context of endogenous *ATM*. **F**. The splicing reporters with *ATM* exon 40 was transfected into HeLa cells and splicing was evaluated with RT-PCR.

We cloned *BRCA2* exon 3 and *ATM* exon 40 into the pSPL3 splicing reporter and introduced selected mutations. Next we also subcloned the WT and mutated exons into the *ACADM* minigene used in our previous study (17), substituting the vulnerable *ACADM* exon 5. We inserted *BRCA2* exon 3 and *ATM* exon 40 into both the WT *ACADM* minigene context, as well as into a context where the strength of the downstream 3’ss is increased from 4.77 (MaxEnt score) to 8.63 (MaxEnt score) in order to test the effect of increasing vulnerability by altering the context of the inserted exons (figure 5C+E). Interestingly, we observed that when the vulnerable *BRCA2* exon 3 is inserted into the pSPL3 splicing reporter, both the c.100G>A and the c.145G>T mutation causes some, but not total exon skipping. The c. 145G>T mutation had the strongest effect on splicing, possibly due to the hnRNPA1-binding ESS, which is no longer inhibited by SRSF1 binding (figure 5D). Surprisingly, when we inserted the vulnerable *BRCA2* exon 3 into the context of the vulnerable *ACADM* exon 5, neither of the two mutations were able to cause exon skipping. When the downstream 3’ss was strengthened, the more severe c.145G>T mutation was able to cause a low degree of exon 3 skipping, but not to the same extent as in the pSPL3 splicing reporter, which has a lower 3’ss score and a longer downstream intron (figure 5D). This indicates, that whereas *BRCA2* exon 3 may be vulnerable in its endogenous context and in the pSPL3 splicing reporter context, the flanking exons of *ACADM* exon 5 do not yield sufficient competition, and the exon becomes resilient in the more favorable *ACADM* exon 5 minigene context.

When we inserted the resilient *ATM* exon 40 into the pSPL3 splicing reporter, we surprisingly observed exon skipping from both the natural c.5932G>T mutation, as well as from the weaker artificial c.5935G>A mutation (figure 5F). As expected, the c.5932G>T mutation had a much more severe effect on splicing, causing almost complete exon skipping, but the c.5935G>A mutation was also able to cause partial exon skipping. When we inserted *ATM* exon 40 into the *ACADM* exon 5 minigene context, we observed that only the more severe c.5932G>T mutation was able to cause exon skipping, but when the downstream 3’ss was optimized, we again observed exon skipping from both mutations, with the strongest effect caused by the c.5932G>T mutation. This illustrates, that the resilient *ATM* exon 40 is vulnerable in the pSPL3 context, resilient in the WT *ACADM* exon 5 context, and vulnerable again in the optimized *ACADM* contexts. Aside from the downstream 3’ss strength, a major difference between the pSPL3 and *ACADM* minigene is the length of the downstream intron. Interestingly, the downstream intron length is the only significant difference between the vulnerable and resilient exons in the ESM database (supplementary figure S8), with vulnerable exons generally having longer downstream introns.

Taken together these results suggest that although *ATM* exon 40 is scored as resilient in the endogenous context and *BRCA2* exon 3 is scored as vulnerable in the endogenous context they behave differently when inserted in the splicing reporters and minigenes. This underscores the importance of assessing exon vulnerability or resilience to splicing mutations in the endogenous context. Since VulExMap is based on empirical data on exon skipping levels from RNA sequencing data from an endogenous context, we believe that it will be a useful when assessing the effect of exonic splicing mutations.

## Discussion

Based on our studies of the vulnerable exon 5 in *ACADM* (17), we hypothesized that the inherent vulnerability to ESMs is different between exons and that this vulnerability is reflected in the observed low levels of basal exon skipping of an exon in the endogenous context in normal cells (27,28). Consistent with this hypothesis, it has also been reported from other genes, like *HRAS* (47), *BRCA2* (29,30) and *CFTR* (31,32,50), that some wild type constitutive exons where ESMs have been reported are slightly less efficiently included in normal tissues. We therefore speculated if this is a general feature of vulnerable exons, which could be used to pinpoint vulnerability to ESMs. Consequently, we designed VulExMap, which identifies low levels of exon skipping in RNA-seq data and we demonstrate in the present study that this approach enables easy identification of vulnerable exons. We previously also observed that *ACADM* exon 2 is skipped in a small proportion of cDNA from patient and control cells (27,28) and consistent with this it was scored as vulnerable by VulExMap analysis. Here we confirm that several mutations in *ACADM* exon 2 associated with MCADD affected splicing in a splicing reporter and that this mimicked ESM-based exon 2 skipping observed in patient cells (figure 1D and Supplementary figure S3). We previously observed the phenomenon of low levels of exon skipping of *HRAS* exon 2 in normal cells, where an exonic mutation in a patient with Costello syndrome was predicted to cause the severe p.Gly12Val change, but instead functions as an ESM that causes a milder disease phenotype (47). VulExMap also confirmed that *HRAS* exon 2 is vulnerable (supplementary figure S9, table S1).

With 45 reported ESMs, *BRCA2* is by far the gene with the highest number of reported ESMs (supplementary table S1). These are clustered in 7 exons (exon 3, 5, 7, 12, 18, 19 and 23 NM_000059), of which all, except exon 23 (containing only one ESM) are predicted by VulExMap to be vulnerable (figure 6A). A recent study of 50 *BRCA2* exon 3 variants revealed that 30 resulted in some degree of exon 3 skipping (30) and another study reported that 32 of 52 selected exonic variants in exon 17 and exon 18 caused exon skipping (29).

**Figure 6.**
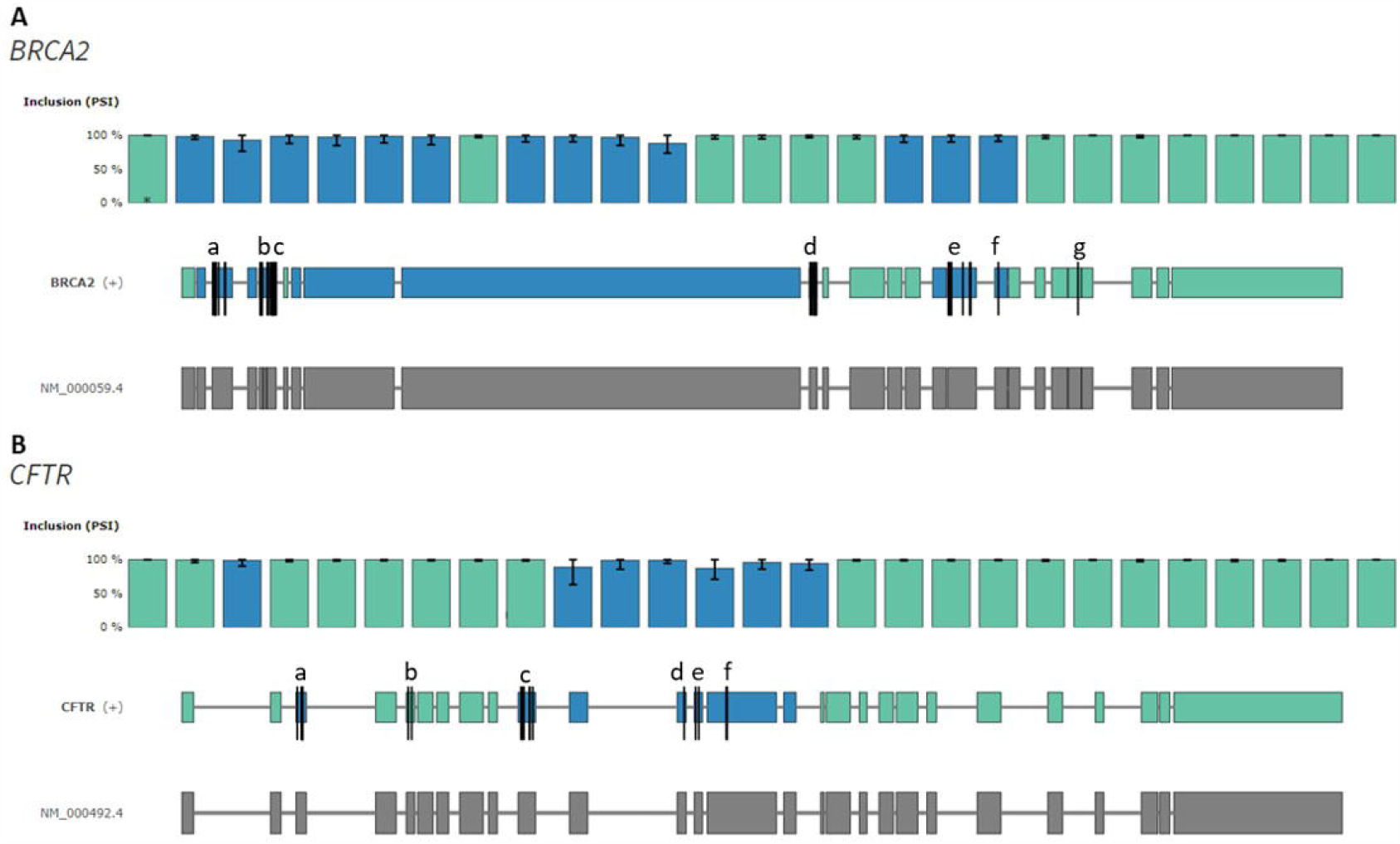
VulExMap shows enrichment of ESMs in known vulnerable exons in *BRCA2* and *CFTR*. **A**. VulExMap of *BRCA2* with GTEx data. Black lines indicate ESMs; 9 in the vulnerable exon 3 (a), 5 in the vulnerable exon 5 (b), 11 I the vulnerable exon 7 (c), 8 in the vulnerable exon 12 (d), 9 in the vulnerable exon 18 (e), 1 in the vulnerable exon 19 (f), and 1 in the resilient exon 24 (g). **B**. VulExMap of *CFTR* with GTEx data. Black line indicates ESMs; 5 in the vulnerable exon 3 (a), 2 in the resilient exon 5 (b), 6 in the vulnerable exon 10 (c), 1 in the vulnerable exon 12 (d), 2 in the vulnerable exon 13 (e), and 2 in the vulnerable exon 14 (f).

In *CFTR*, five of the six exons where ESMs have been reported (exon 3, 5, 10, 12, 13 and 14, NM_000492) are vulnerable according to VulExMap (figure 6B) and only one (exon 5) is resilient. Consistent with this, low levels of exon skipping from normal cells have been reported from *CFTR* exons 10, 13 and 14 (31,51,52). ESMs have been reported based on initial analysis of patient mRNA in the predicted vulnerable *CFTR* exons 3, 10, 12, 13, and 14, whereas the two reported ESMs in the resilient exon 5 have only been analyzed in a hybrid minigene (32), where the strength of the upstream 5’ss was increased. This would alter the vulnerability of *CFTR* exon 5 in the reporter and could thereby lead to artificially high exon skipping from the two reported ESMs. We show in the present study that vulnerability of *BRCA2* exon 3 and *ATM* exon 40 to ESMs is context-specific and we have previously shown that the splice sites of neighboring exons and intron size are important in defining vulnerability by analyzing splicing of WT and mutant *ACADM* exon 5 and exon 6 in different contexts (17). Consistent with our studies, Hefferon and co-workers (53), who analyzed *CFTR* exon 10 splicing kinetics, showed that also strengthening of the upstream (exon 9) 5’ss weakens definition of the downstream exon and therefore increases vulnerability. Moreover, a recent study (54) also demonstrated that the strength of the upstream 5’ss has a strong effect on inclusion of the downstream exons employing a massive parallel splicing assay. This is consistent with our observations on the splicing patterns observed when *BRCA2* exon 3 and *ATM* exon 40 are inserted into different contexts and suggests that using splicing reporters, such as the pSPL3 vector, may not always accurately mimic the effect a mutation may have on splicing.

At present, exons in a disease gene are considered equally likely to be affected by ESMs and the proportion of exonic mutations that cause exon skipping by affecting splicing regulatory elements (SREs) has been reported to be as high as 77% in *MLH1* exon 10 and 60% in *BRCA2* exon 3 (30,55). Our analysis with VulExMap demonstrate that both *MLH1* exon 10 and *BRCA2* exon 3 are vulnerable (supplementary table S1) and therefore more sensitive to ESMs. On the other hand, Soemedi et al. tested 4,964 exonic disease associated mutations in a fixed genomic context employing a splicing reporter and found that only 10% of the mutations altered splicing (16). However, the splicing reporter in their assay consisted of a design with an atypical context, where only exons ≤100 bp was used and the flanking introns were only ∼200-300 bp long (supplementary figure S10). This is far below the mean length of exons and flanking introns (mean exon length is ∼150 bp, mean upstream intron length is ∼7000 bp, and mean downstream intron length is ∼6000 bp) and may have resulted in a very efficient splicing reporter, which is only sensitive to mutations that cause a major alteration to the splicing code. Furthermore, as we have observed that downstream intron length is directly corelated with exon vulnerability, the very short downstream intron may increase the splicing efficiency, and thus resilience of the investigated exons. On average the group of vulnerable exons identified by VulExMap are more likely to have a weaker 3’ss than that of the flanking downstream exon (figure 2F-G). This may contribute to explain why these exons are particularly vulnerable to ESMs as it is well-documented that the cooperation and competition of the up- and downstream splice sites influence the efficiency of exon inclusion (56-58). This further underscores that when testing the effects of exonic mutations on splicing by employing minigenes, it is imperative that these very closely reflect the endogenous context.

Although the vast majority (72%) of reported ESMs are located in vulnerable exons there is, as mentioned above, also a proportion of ESMs, which are located in resilient exons (supplementary table S1). Although some of these ESMs may have been misclassified because exon skipping has only been observed by minigene analysis and not in the endogenous context, it is also apparent that disrupted splicing is not simply an all or none process. Our previous analysis of *ACADM* exon 5 mutations (17), as well as the present analysis of mutations in *BRCA2* exon 3 and *ATM* exon 40 indicate that the effect a mutation has on splicing is highly dependent both on the preexisting balance between ESEs and ESSs in the affected exon and the alteration (ESE loss or ESS gain vs ESE loss + ESS gain) to the splicing code caused by the mutation. The TAGACA hnRNPA1 binding motif causes exon skipping when created in *ACADM* exon 2, *F9* exon 5 and *BRCA2* exon 12 (59,60) (figure 7A), which are all vulnerable. When this motif is created by a mutation in the resilient *CDC73* exon 2 it does not cause exon skipping (4), suggesting that simple creation of this ESS motif is not sufficient to cause exon skipping of a resilient exon. However, a mutation that also creates the TAGACAA motif does cause skipping when created in the resilient *HPRT1* exon 8. Interestingly, the mutation in *HPRT* exon 8 simultaneously abolishes an SRSF1-binding ESE (CAGACAA), identical to that in *SMN1* exon 7. The ESM in the resilient *HPRT* exon 8 therefore causes an identical dual event to the simultaneous loss of an ESE and gain of an ESS documented to cause *SMN2* Exon 7 skipping, where the SRSF1-binding ESE, CAGACAA (*SMN1*), is altered to the TAGACAA hnRNPA1-binding ESS (*SMN2*). This indicates that skipping of a resilient exon requires a dual event. It reflects that the relative importance of the affected SRE is determined by exon vulnerability, but that also the severity of the change that the mutation imposes on the SRE is important (i.e. whether skipping can be caused by creating an ESS *or* by disrupting an ESE (single change to the splicing code), or if the ESM simultaneously need to create an ESS *and* disrupt an ESE (dual change to the splicing code) (figure 7B). Our PESE and PESE analysis (figure 2A+B) revealed that resilient exons are more strongly defined, due to the overrepresentation of PESEs and underrepresentation of PESSs compared to vulnerable exons. This is consistent with recent findings (15), where it was demonstrated that alternative exons are enriched with suboptimal ESEs, that are only one mutation away from becoming an ESS (reference).

**Figure 7.**
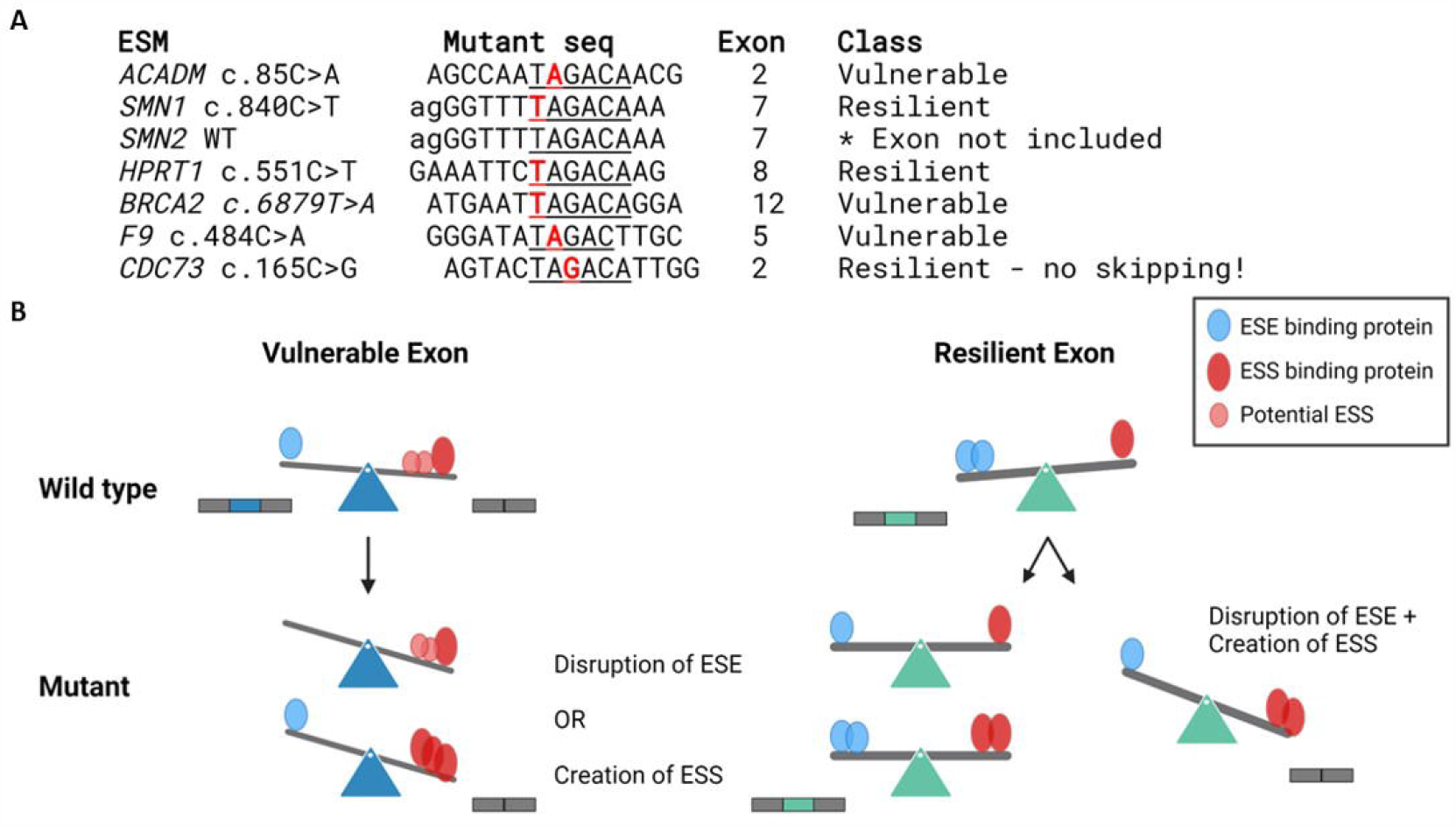
Mutations creating the same motif does not always result in aberrant splicing. **A**. The TAGACA hnRNPA1 ESS created by the c.85C>A in *ACADM* exon 2, is also created in the resilient *SMN1* exon 7 (c.84C>T) and is the primary reason why *SMN2* exon 7 is not included in the final *SMN2* transcript. This ESS is also created in the resilient *HPRT1* exon 8 (c.551C>T), and in the vulnerable *BRCA2* exon 12 (c.6879T>A). A TAGAC motif in also created in the vulnerable *F9* exon 5 (c.484C>A), but in the resilient *CDC73* exon 2 (c.165C>T) creation of the full TAGACA motif does not result in exon skipping, indication that this resilient exon is too strongly defined to be skipped by creation of a single ESS but would probably require a dual change to the splicing code. **B**. Model of single and dual change to the splicing code. When a mutation disrupts an ESE or creates an ESS, the potential ESS elements across the vulnerable exon will attract a negative splicing factor and cause exon skipping by displacing the positive splicing factors. Resilient exons have a higher density of ESEs, as well as stronger splice sites, and are therefore better suited to handle single changes to the splicing code. A mutation that only disrupts an ESE or creates an ESS will often not be sufficient to cause exon skipping. Only if a mutation severely alters the splicing code, by both disrupting an ESE *and* creating an ESS (dual change), will the balance be shifted sufficiently to cause skipping of the exon.

Previous studies investigating the predictive power of *in silico* tools, such as ΔtESRseq (61) and ΔΨ (5), found that the accuracy of these tools varies for different genes and exons (55). Because they simply score if an altered motif results from the mutation, and this is more likely to affect a vulnerable exon. Consistent with our findings, Canson et al. recently reported that these tools have the highest accuracy in *BRCA1* exon 6, *BRCA2* exon 7, *CFTR* exon 12, *MLH1* exon 10 and *NF1* exon 37 (62), which are all vulnerable according to VulExMap analysis (supplementary table S1). Recent studies have implemented new decision pipelines for determining pathogenicity of exonic variants (30,63), and we suggest that knowledge of vulnerability should also be included to aid such classification. While tools such as HexoSplice (13) and EX-SKIP (4) were able to predict a splicing change in the majority of the mutations in the ESM database, there was no significant difference in the predicted scores between resilient and vulnerable exons (supplementary figure S11).

In summary, we have developed VulExMap, which based on empirical data, classifies the vulnerability of an exon, when present in its endogenous genomic context. We used it to demonstrate that, on a transcriptome wide basis, vulnerability to ESMs differs between constitutive exons. Importantly, VulExMap provides a simplistic and user-friendly way to evaluate whether a mutation is located in a vulnerable exon and therefore has a several fold higher risk of causing effects on splicing. Therefore, we believe that VulExMap will be useful in future genetic diagnosis.

## Acknowledgements

We thank Charles Stanley, Childrens Hospital of Philadelphia, for providing material and info about MCADD patients. We thank Tine Christensen, Mette Larsen, Kasper W. Stonor, Tanja Bruun, Kristian Traantoft Rasmussen, Margrethe Thusholdt, Krystyna Giemza and Aleksandra Kulus for technical assistance.

