## Supplemental figures for "All exons are not created equal - Exon vulnerability determines the effect of exonic mutations on splicing"

### Supplementary figures

#### Supplementary figure S1

Overview of the VulExMap classification scheme. Splice juncion counts from Snaptron are pulled from the webserver for regions covering each gene in the Refseq annotation and splice junctions with one or more unannotated splice-sites are filtered out to remove potentially false mappings. The junction counts are then used to obtain percent-spliced-in (PSI) estimates for each sample and each type of alternative splicing pattern and these estimates are then used to compute the coefficient of variation (CV) for cassette exons. Only samples with a sum of splice jucntion counts of at least 10 for a given splicing pattern is used to compute mean PSI and CV. Cassette exons with a mean splice junction count per sample of at least 30 are then binned into PSI classes based on the mean PSI of the exon, and vulnerability thresholds are determined using the median of the 80-85% PSI bin and the median of the 99-99.5% bin. Following determination of these thresholds, classification of exonic segments is applied broadly to all segments. An exon or an exonic segment is considered resilient if the mean PSI is at least 99.5% and the CV is below the minimum threshold determined for vulnerability. If the mean PSI is at least 85% and the CV is lower than the upper CV threshold limit it is considered vulnerable. If it has a mean PSI below 85%, or a CV higher than the upper threshold for vulnerability, it is considered alternative.

#### Supplementary figure S2

Screenshots from vulexmap.compbio.sdu.dk. **A**. VulExMap front page. **B**. Gene table where the user can select the gene of interest for plotting. **C**. VulExMap plot of *ACADM*, The bar chart show PSI for each segment, the error bars represent the 95% inter-quantile range. Bellow the bar chart a gene model is made with all segments colored by classification. Bellow the genemodel are all refseq transcripts for the chosen gene. The top transcripts, NM_000016.4 represent the canonical protein coding transcript. **D**. When hovering above the bar chart, the PSI, SD, 95%IQR and CV can be seen for the chosen segments, along with information about how many samples have been included or excluded from the analyses, and the mean reads. **E**. When clicking on a bar the raw inclusion reads for the chosen segment is shown. **F**. The ESMs from this paper is already included in VulExMap. The user can zoom in on the gene model and hover the cursor above the black lines to get information about the ESM. **G**. The user can upload their own list of mutations in a bed6 file to VulExMap, to observe the location of the mutations. **H.** The uploaded mutations will then be observed on the gene model. If a mutation is located in the first or last 3 bp of an exon, it will be colored red. This is to indicate that the mutation is located in the splice site, and that any effect on splicing is independent of vulnerability.

#### Supplementary figure S3

Confirmation of the *ACADM* exon 2 splicing defect in patient cells. A MCADD patient with the common c.985A>G mutation on one allele and an unknown silent c.87A>G mutation on the other allele was investigated for splicing mutations. **A**. Cells from two control, the patient, and their father, both heterozygous for c.87A>G, were cultured with and without cycloheximide (CHX). Increased skipped product in the +CHX samples from the patient and their father indicates that c.87A>G causes *ACADM* exon 2 skipping and NMD in patient cells. **B.** Sequence analysis of patient cDNA confirms that the patient is heterozygous for the c.87A>G mutation, and that the G allele is dramatically reduced in the full length (exon 2 included ) product.

#### Supplementary figure S4

dbSNP156 variants and sQTL distribution across vulnerable, resilient, and alternative exons. **A**. distribution of dbSNP156 variants/nt of the three exon classes. **B**. distribution of sQTLs/nt of the three exon classes. **C**. distribution of synonymous variants/nt of the three exons classes. **D**. distribution of missense variants/nt of the three exon classes. **E**. ratio between synonymous and missense variants in the three classes.

**Supplementary figure S5**

K-mer scatter plots between the resilient vs. vulnerable, resilient vs alternative, and vulnerable vs. alternative exons. While the are very closely related with only few significantly enriched k-mers, the resilient vs. alternative exons have a higher dispersion and more enriched k-mers that a different between the two groups. The vulnerable and alternative k-mes are closer than the resilient vs. alternatve, but more diverse than the resilient vs. vulnerable.

**Supplementary figure S6**

DeepCLIP predictions of SRSF1 and hnRNPA1 binding to the two *BRCA2* mutations, c.100G>A and c.145G>T. the data used for the predictions was for SRSF1 ENCODE eCLIP data from K562 (1), and for hnRNPA1 iCLIP from Bruun et al. 2016 (2).

**Supplementary figure S7**

DeepCLIP predictions of SRSF1 and hnRNPA1 binding to the two *ATM* mutations, c.5932G>T and c.5935G>A. The data used for the predictions was for SRSF1 ENCODE eCLIP data from K562 (1), and for hnRNPA1 iCLIP from Bruun et al. 2016 (2).

**Supplementary figure S8**

Resilient and vulnerable exons from the ESM database were filtered so each exon was only represented once. The resilient (n = 61) and vulnerable (n = 82) were analyzed for characteristics of exon definition; PESE content, PESS content, GC content, Exon length, upstream intron length, downstream intron length, 3’ss strength, 5’ss strength, 5’ss difference from upstream 5’ss and 3’s difference from downstream 3’ss. Wilcoxon rank sum test was performed between the two groups, *** $p < 0.0005$.

**Supplementary figure S9**

VulExMap of *HRAS.* Shown the black line indicates a c.35GC>TG mutation in *HRAS* the vulnerable exon 2 which causes exon skipping (3).

**Supplementary figure S10**

**A.** Comparison between the splicing reporters used in this study and the MaPSy vector from Soemedi et al. (4). **B**. Mean values for splice site score and exon/intron lengths for all exons in the investigated GTEx data, after filtering.

**Supplementary figure S11**

Ex-SKIP prediction (5) and Hexosplice prediction (6) could not distinguish between the mutations in vulnerable or resilient exons.

**Supplementary figure S1**

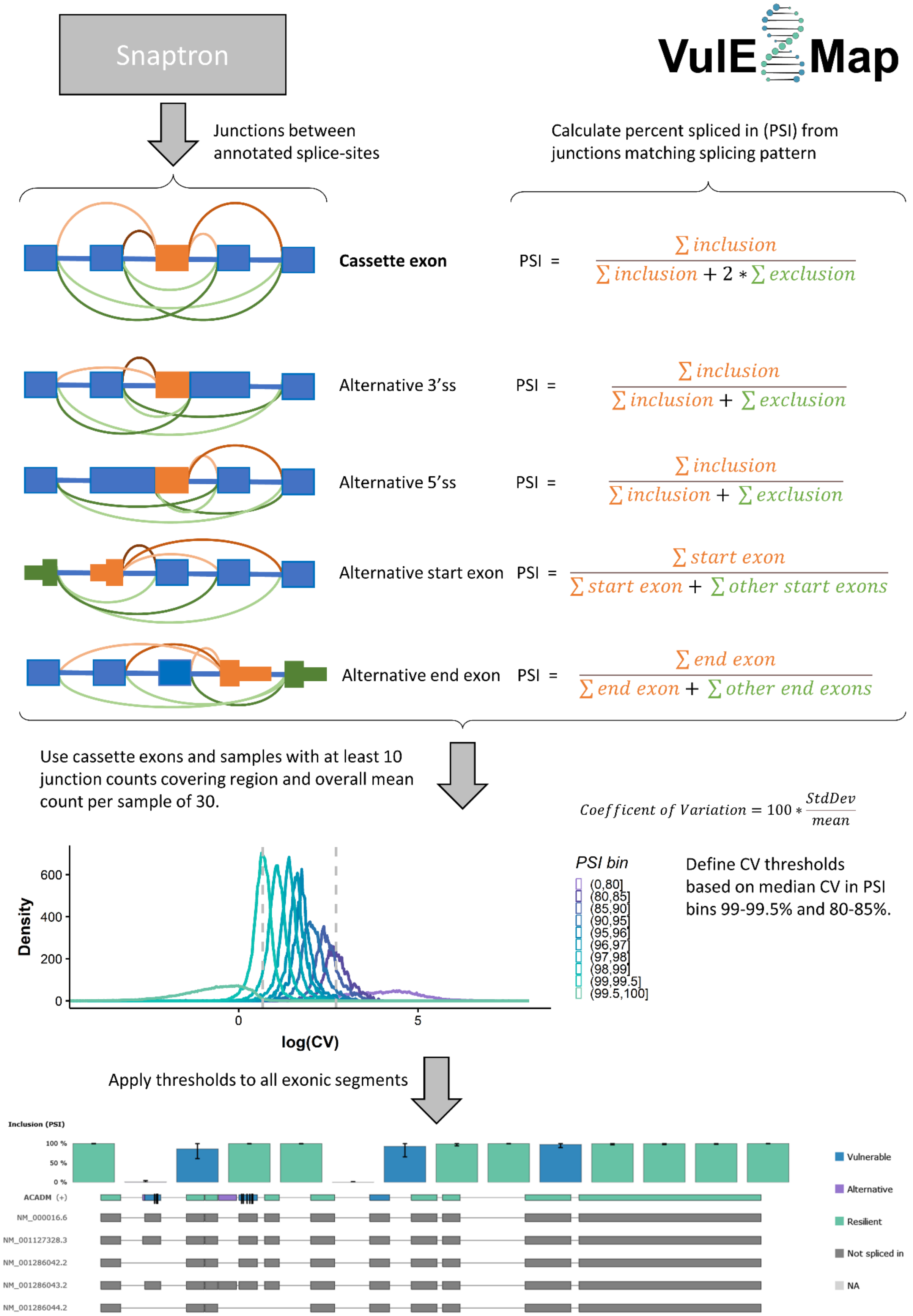

**Supplementary figure S2**

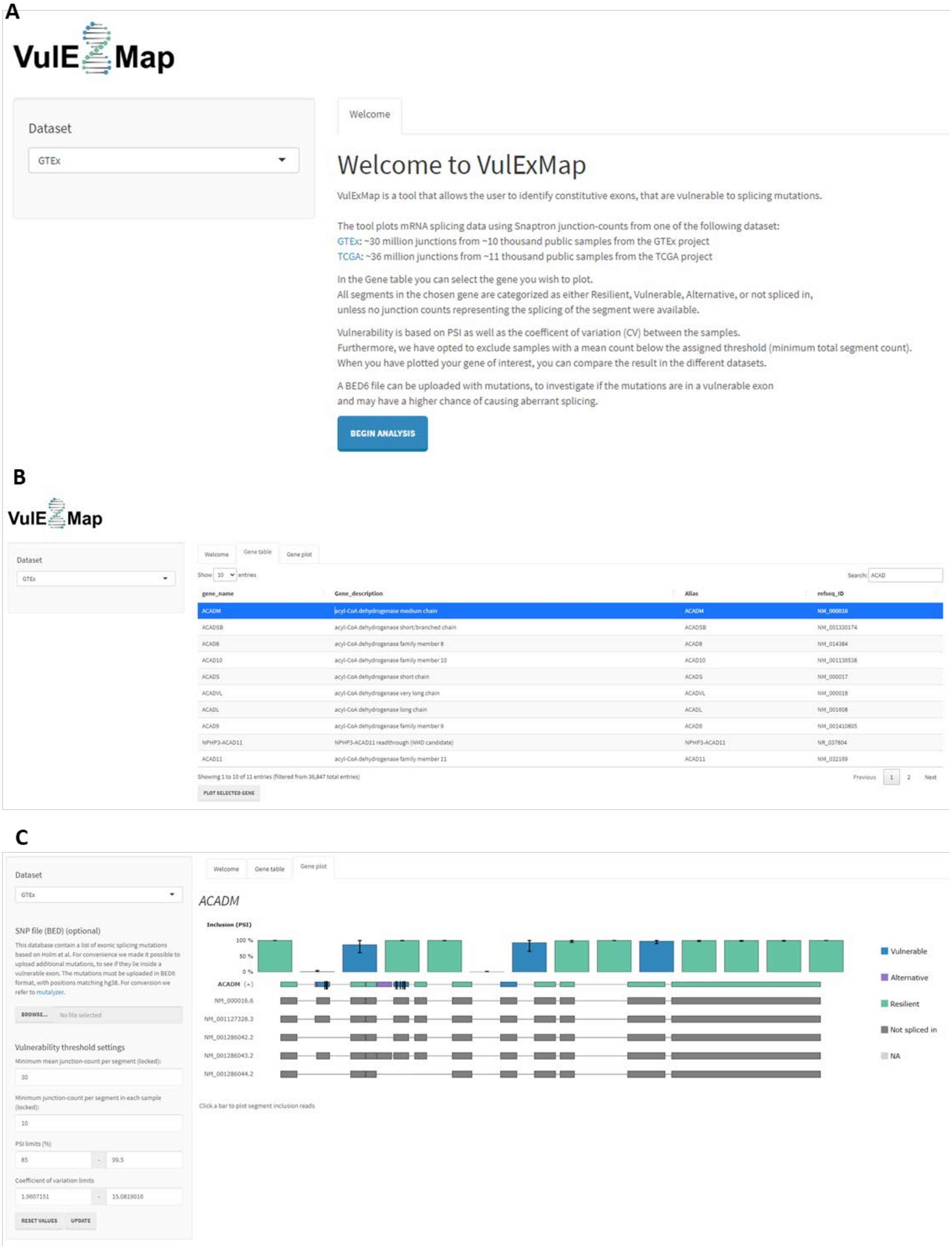

Supplementary figures. Holm et al. 2023

#### Supplementary figure S2

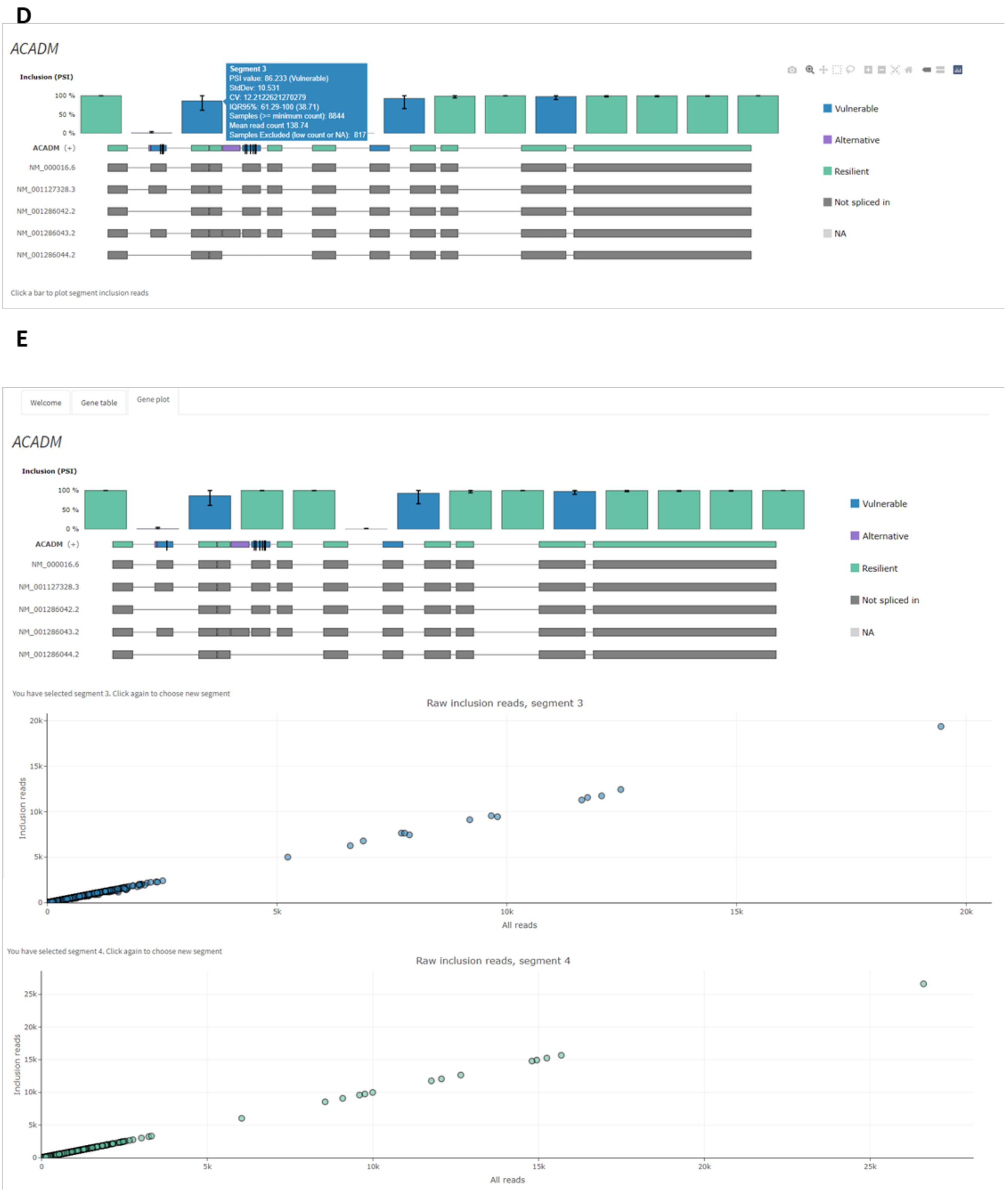

Supplementary figures. Holm et al. 2023

**Supplementary figure S2**

**F**

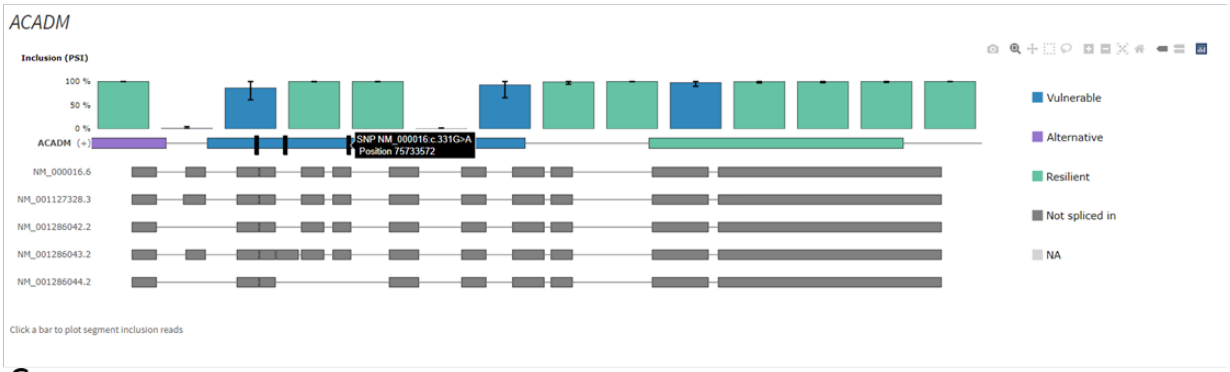


**G**

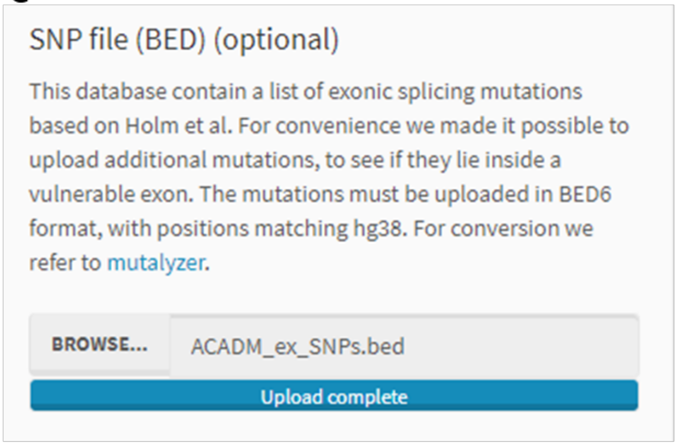


**H**

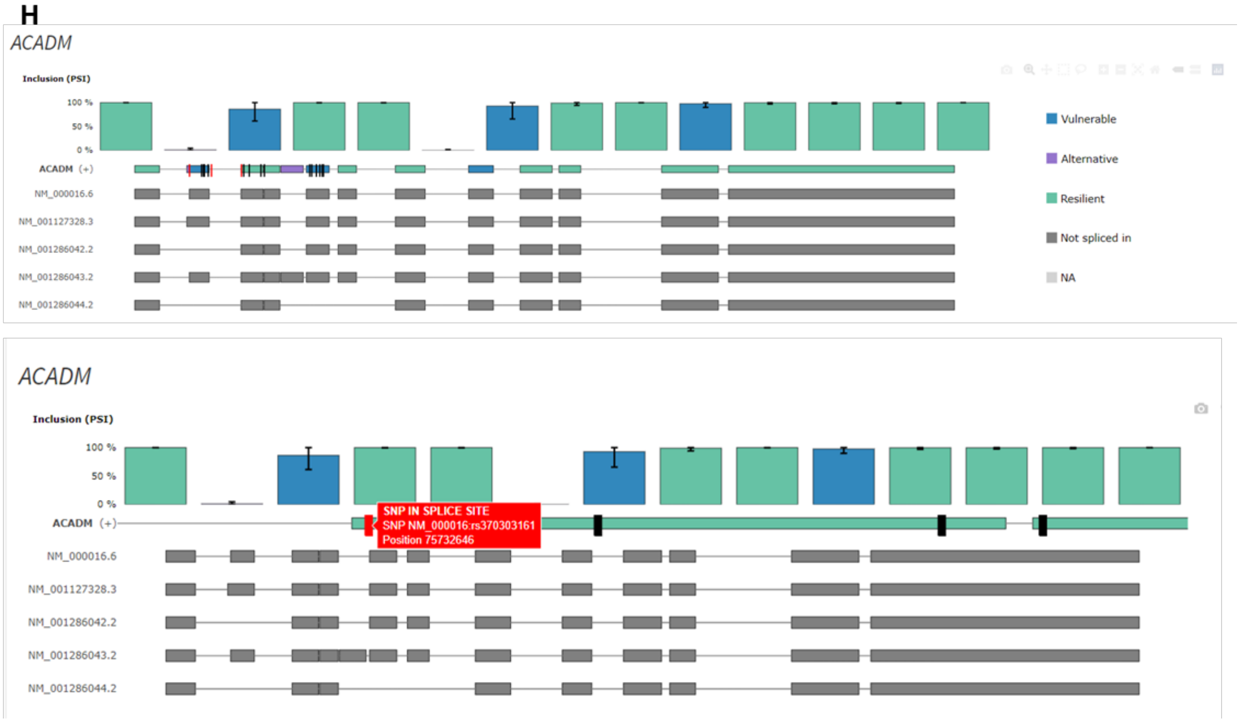

**Supplementary figure S3**

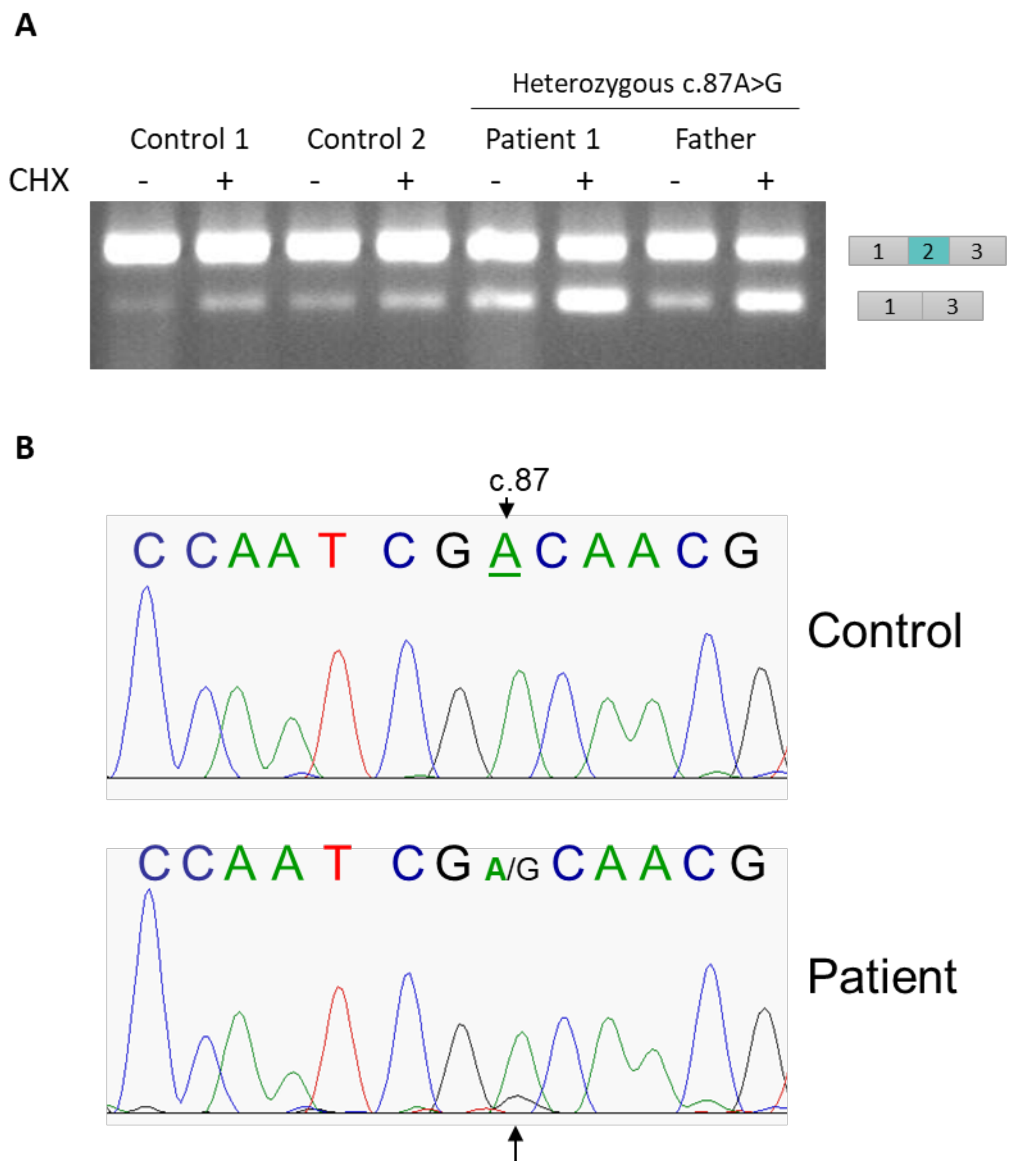

#### Supplementary figure S4

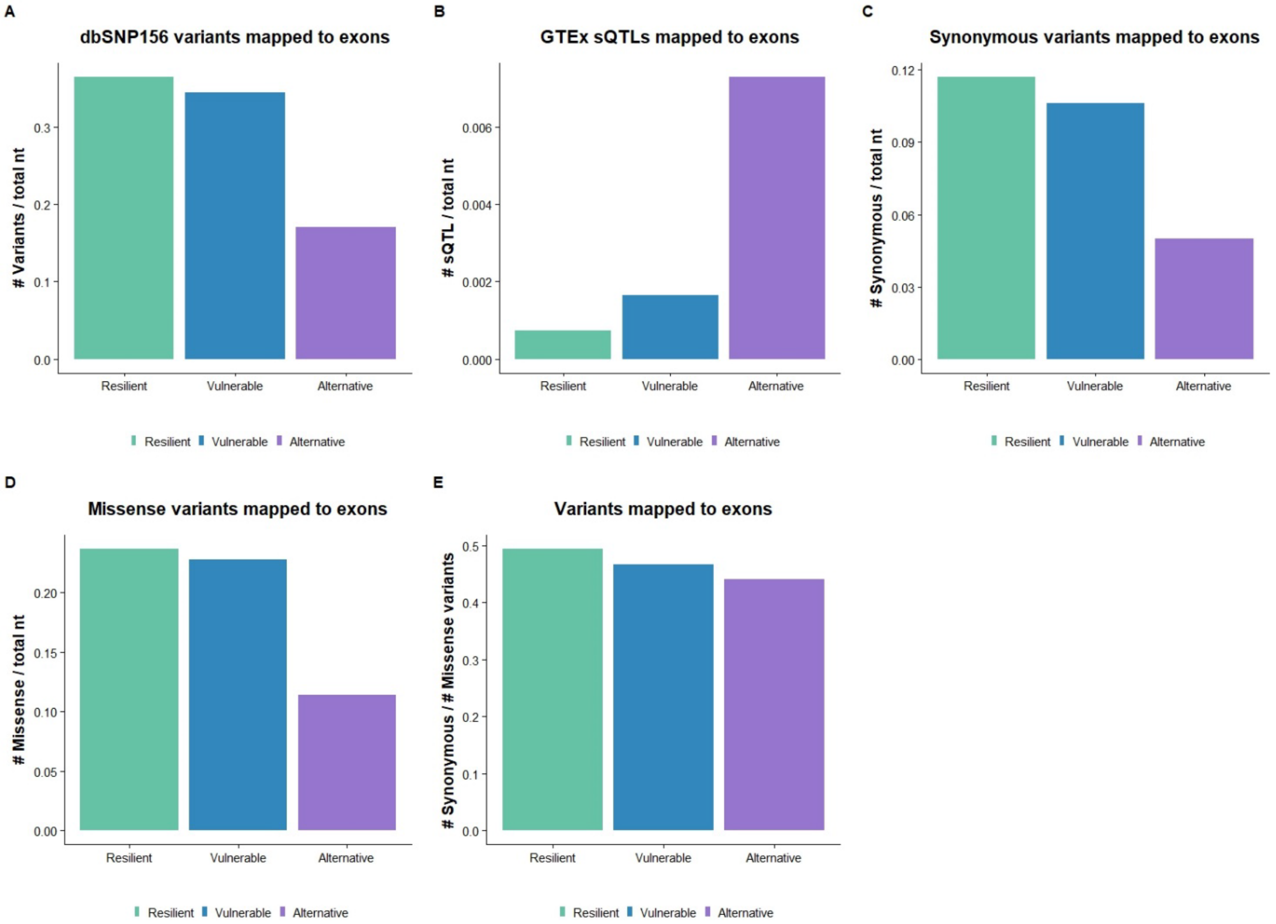

**Supplementary figure S5**

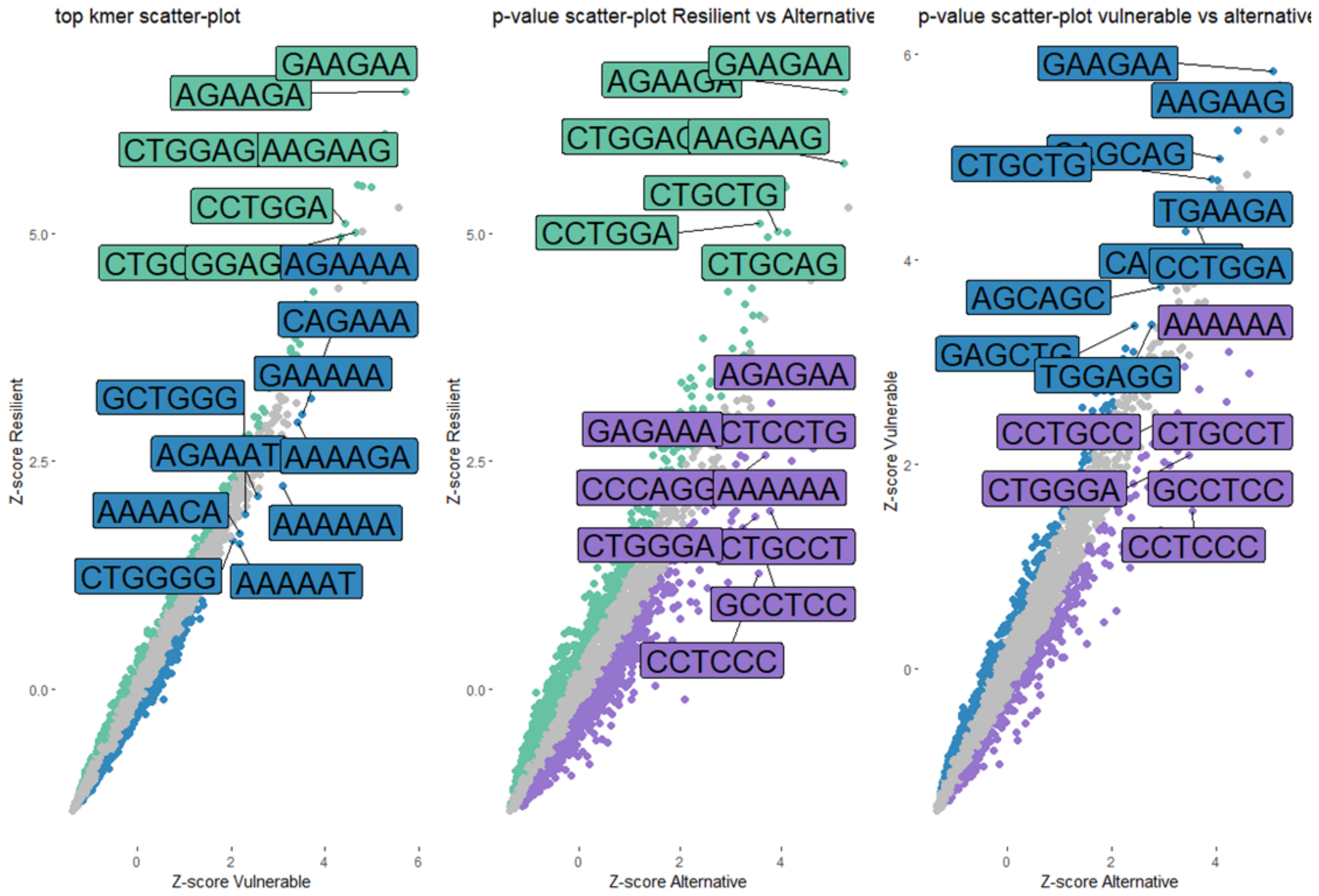

**Supplementary figure S6**

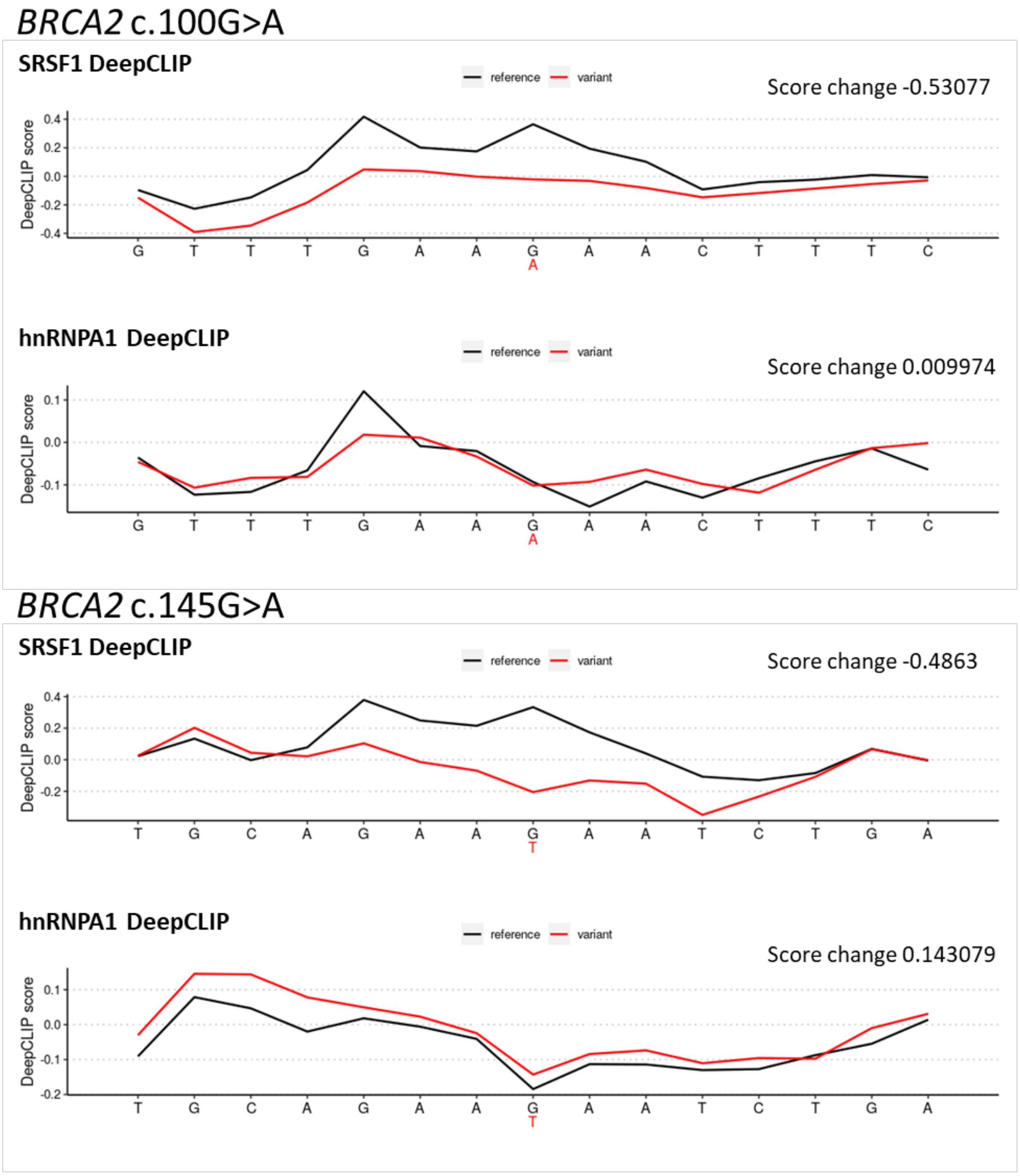


| c.100G>A | c.145G>T |
|---|---|
| SRSF1 ↓ | SRSF1 ↓ |
| hnRNPA1 | hnRNPA1 ↑ |

**Supplementary figure S7**

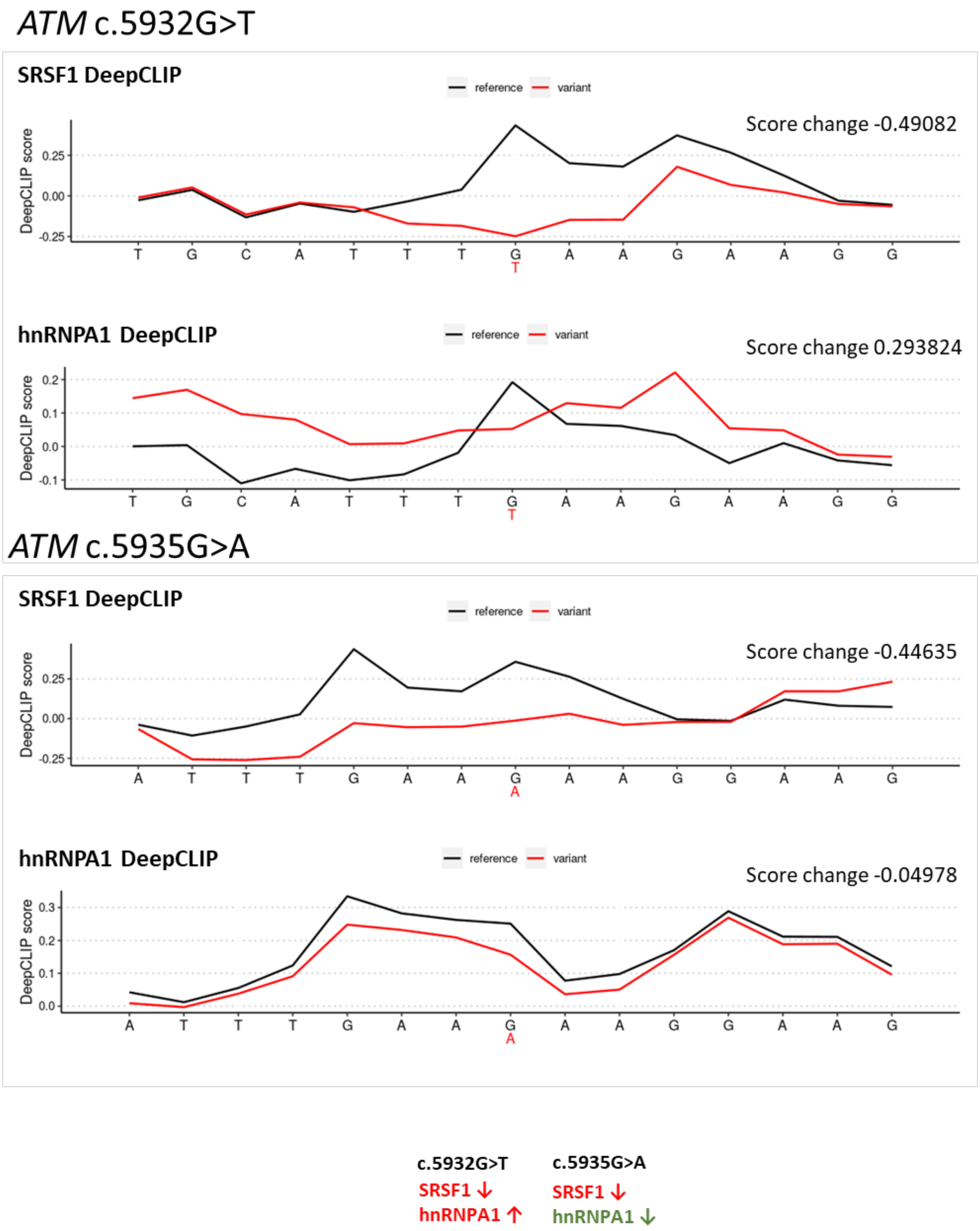

#### Supplementary figure S8

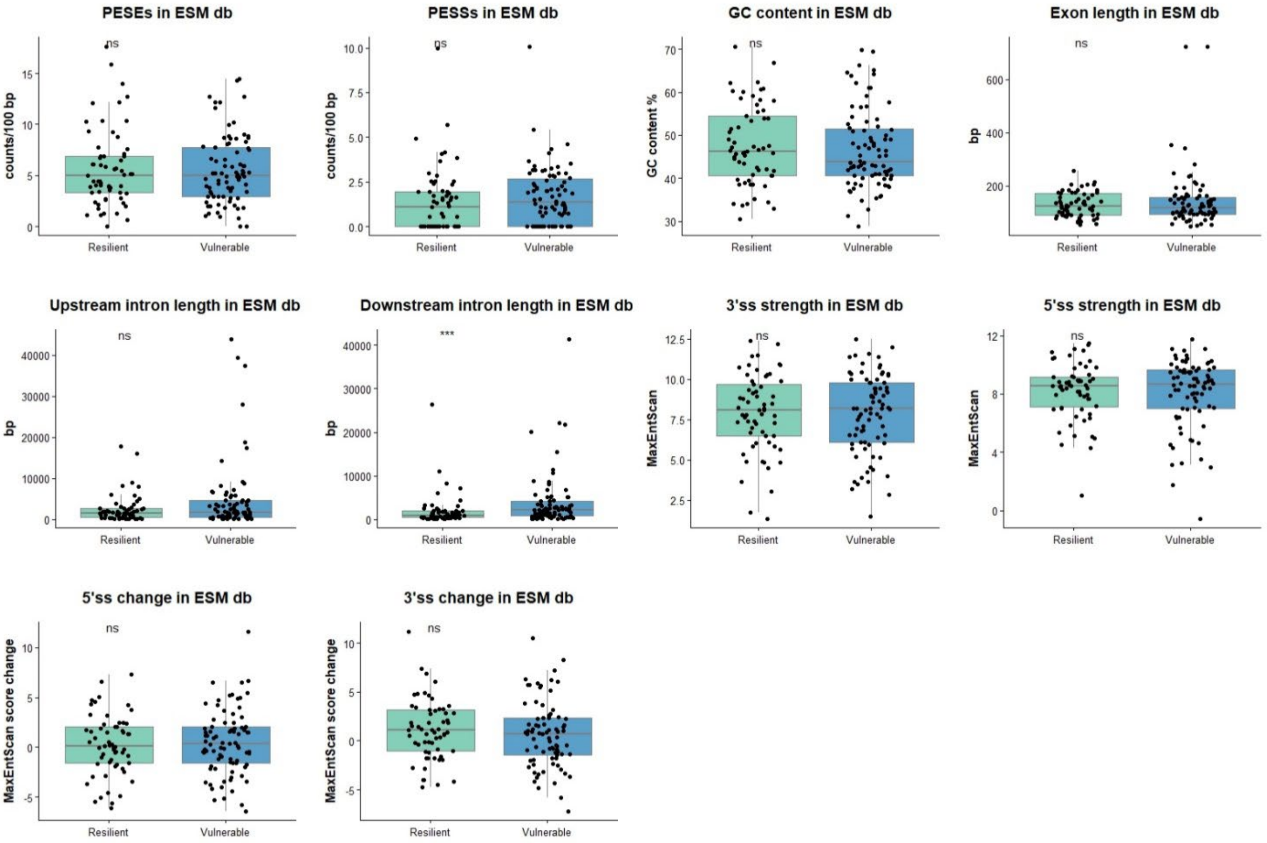

#### Supplementary figure S9

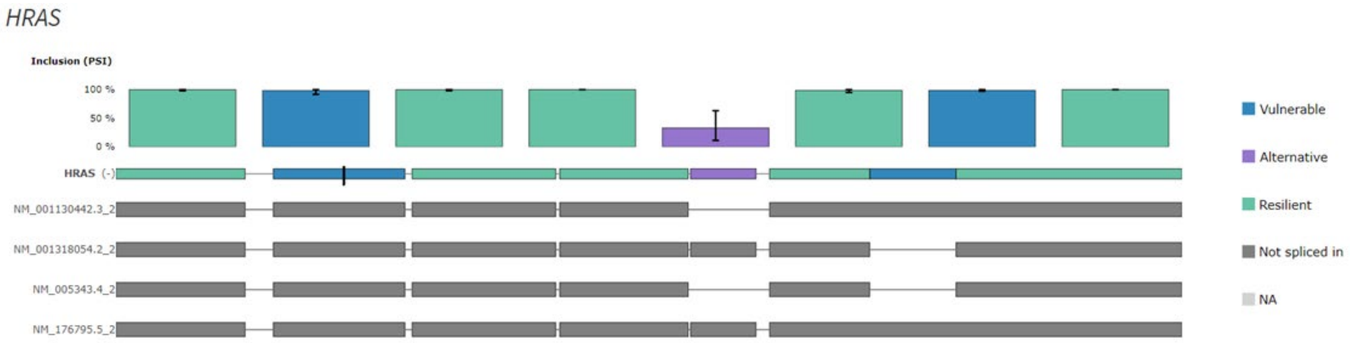

**Supplementary figure S10**

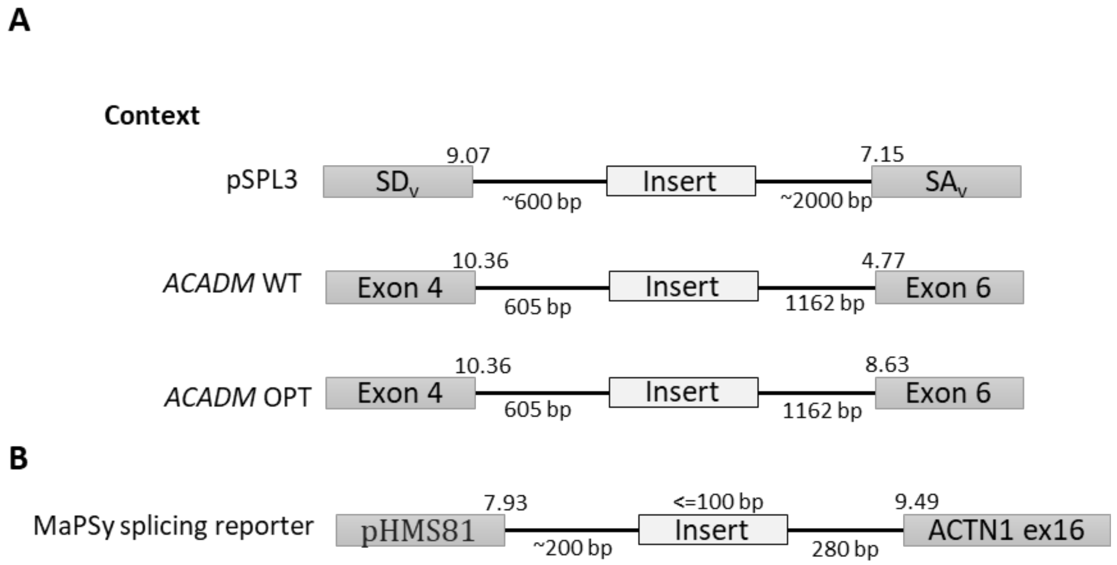


Mean values for all exons:

| | |
|---|---|
| 3’ss score | 8.22 MaxEntScan |
| 5’ss score | 8.11 MaxEntScan |
| Exon length | 154 bp |
| Upstream intron Length | 7257 bp |
| Downstream intron length | 6374 bp |

#### Supplementary figure S11

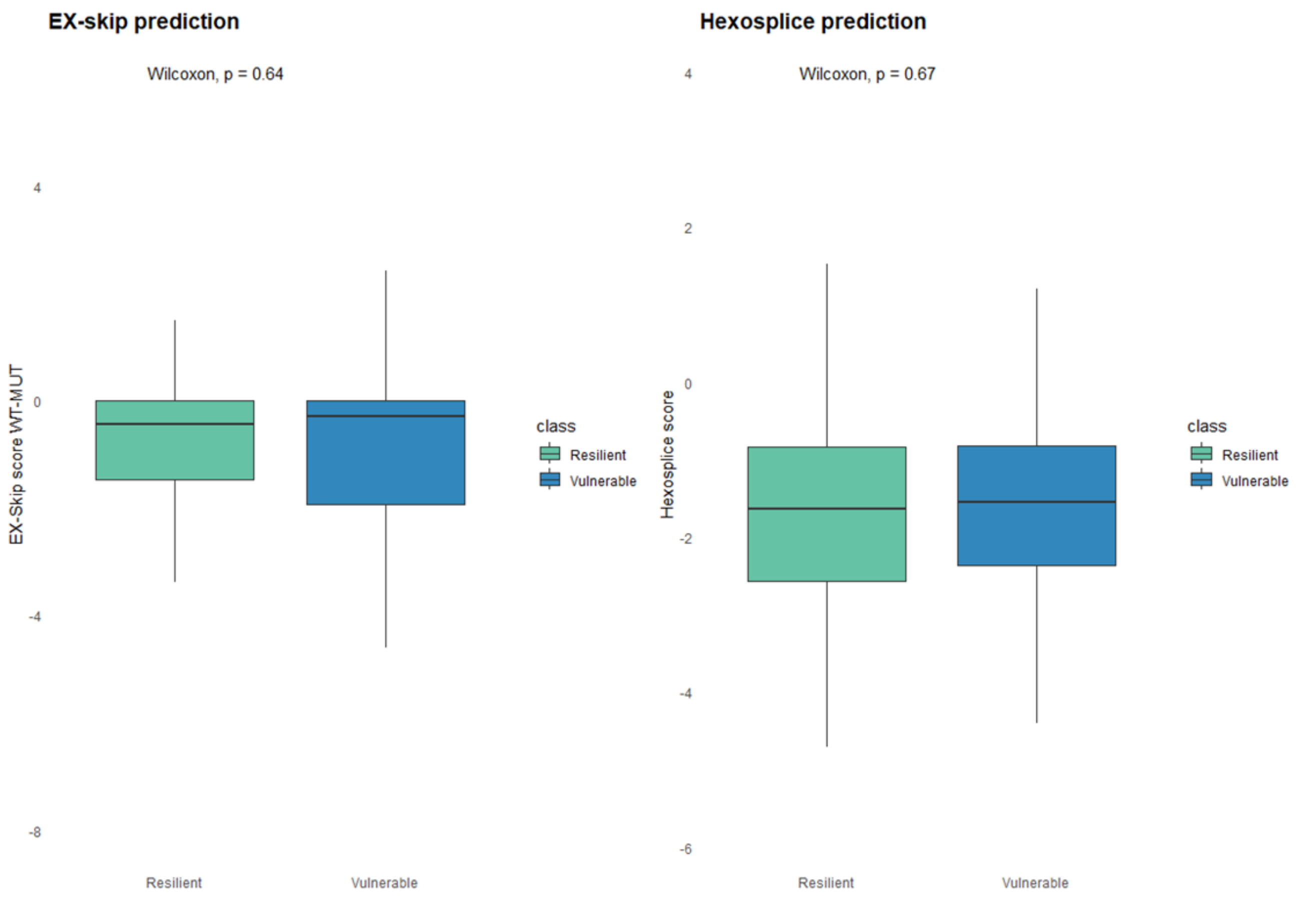
